# Structural patterns and transcriptional effects of integrated Epstein-Barr virus revealed by long-read sequencing and RNA-sequencing in Nasopharyngeal Carcinoma

**DOI:** 10.1101/2025.02.15.638273

**Authors:** Zahra Kardan, Adway Kadam, Annie Wai Yeeng Chai, Sok Ching Cheong, Sampath Kumar Loganathan

**Author notes:** Correspondence: Sampath Kumar Loganathan, Cancer Research Program, RI-MUHC, 1001 Decarie Boulevard, Montreal, QC, Canada H4A 3J1.

## Abstract

Integration of Epstein–Barr virus (EBV) DNA into the human genome is a critical event in nasopharyngeal carcinogenesis. Here, we comprehensively characterize large-scale virus-human integration events in four EBV-positive nasopharyngeal carcinoma (NPC) cell lines and tumors using Nanopore long-read sequencing technology. We identified four distinct integration types, with Type I being particularly notable, characterized by continuous long reads covering nearly the entire EBV genome. Our findings reveal the involvement of multiple integration events in inducing inter-chromosomal translocations, leading to significant genomic disruption through chromosomal rearrangements. Additionally, we explore the relationship between EBV integration sites and structural variations, further supporting the role of EBV integration in driving genomic instability. By integrating RNA-seq data, we demonstrate the potential for EBV integration to disrupt gene expression, highlighting several integration sites within cancer-associated genes such as CD96, ARHGAP27, ASH1L, KDM3B, ZMYM2, and PIK3C2A. Notably, EBV-human fusion events were prevalent in EBV-associated NPCs, including intriguing fusion transcripts such as LRRC8C-RPMS1 and LINC00486-RPMS1, which provide further evidence of the oncogenic potential of EBV integration. Taken together, this study uncovers EBV integration patterns in Nasopharyngeal carcinogenesis using long-read sequencing technology.

## Introduction

Nasopharyngeal carcinoma (NPC) is a type of epithelial carcinoma arising from the nasopharyngeal mucosal lining ^1^. NPC exhibits a unique geographical distribution, being highly endemic in East, Southeast Asia, and North Africa while remaining relatively rare in other parts of the world ^2^. However, NPC imposes a considerable global health burden, as indicated by the International Agency for Research on Cancer (IARC) in 2020, with around 133,354 new cases along with approximately 80,008 fatalities associated with the disease worldwide ^3^. NPC often presents with non-specific symptoms such as nasal congestion and ear infections, leading to delays in diagnosis; >70% of patients with NPC are already at stage III or IV at the time of initial diagnosis^4^. Owing to the deep tumor localization and complex anatomical structure of the tumor site, surgical options are often limited ^5^. The remarkable geographical distribution and challenging diagnosis and treatment approaches associated with NPC have prompted studies into its risk factors. EBV infection, host genetics, and environmental influences have been reported to contribute to NPC development^6^ ^7^ ^8^.

EBV infection, involving over 90% of patients with confirmed non-keratinizing NPC, stands as the primary risk factor ^9^ ^10^. NPC also has the highest EBV RNA burden among all EBV-positive cancers ^11^. EBV establishes Type II latency in NPC, resulting in the accumulation of immune cells and immunosuppression by viral pathogens, which contribute to tumor initiation, growth, and development through EBV latency genes ^12^. EBV latency genes include LMP1, LMP2, EBNA1, BART miRNAs, EBERS, and BARF1. These genes manipulate various cellular pathways, which ultimately leads to increased cell proliferation and the regulation of the host microenvironment, which in turn supports the clonal expansion of EBV-infected pre-invasive epithelial cells ^13^. However, the full spectrum of how EBV integration influences host gene function, cellular processes, and genomic stability has remained largely unexplored.

Next-generation sequencing (NGS) technologies, capable of parallel processing millions to billions of short reads (∼150–300 bp), have revolutionized genomic and biomedical research^14^. Several NGS studies have characterized genome-wide alterations in both viral and host genomes due to EBV integration in NPC and gastric carcinoma^15^ ^16^. These studies have shown that integration sites are dispersed across the human genome and occasionally co-localize with regions containing cancer-associated genes, such as JAK2, PD-L1, and PD-L2 in gastric carcinomas, and PARK2, TNFAIP3, and CDK15 in NPC, suggesting that EBV integration may promote carcinogenesis^15^ ^16^. However, NGS requires higher coverage and generates reads with 150–300 bps length, limiting further characterization of EBV integration to the genome. It only allows detection of integration sites (i.e., breakpoints) between the virus and human genomes rather than complete integration events. In addition, it is challenging to identify long or complex structural variations in the host genome upon EBV integration using the current short-read NGS technologies.

To address these challenges, this study performs a comprehensive characterization of large-range virus-human integration events in four EBV-positive NPC samples using Nanopore long-read sequencing technology to capture the whole length of EBV integration. Combined with the corresponding RNA-seq data, this approach sheds light on the cellular pathways and protein-protein interaction in the presence of EBV. This study also analyzes the relationship between EBV integration, various SVs, and their functional consequences in nasopharyngeal cancer. These findings provide insight into EBV integration and its oncogenic effect on NPC progression.

## Material and methods

### Cell line & sample collection

EBV-positive NPC cell lines (C666-1, NPC43 and C17) were maintained in RPMI1640 media supplemented with 10% FBS, 100 IU Pen/Strep, and 4 μmol/L Y-27632 (except for C666-1 was grown without Y-27632) and were authenticated by short tandem repeat profiling. The resected NPC219 tissue was collected from a consented patient at the time of diagnosis, with Institutional Review Board approval.

### DNA extraction, library construction, and nanopore sequencing

Genomic DNA was extracted from 20 to 60 mg of the tumor tissues or cells using a Nanobind® tissue kit (PacBio). The extracted DNA was processed through a short-read elimination kit (Circulomics, PacBio). The DNA concentration for each sample was measured using Nanodrop 3300 (Thermo Fisher Scientific). Sequencing libraries were constructed using the Oxford Nanopore Technologies (ONT) ligation sequencing kit (SQK-LSK109). Prepared libraries were subsequently sequenced using the PromethION platform and 1D flow cell, containing protein pore R.9.4.1. Nanopore sequencing results were processed using ONT’s Guppy software (v6.2.7) (to convert current intensity values (in fast5 format) into nucleic acid sequences. PycoQC (v.2.5.2) was used to perform quality control of the original fastq data, removing adapters, and short reads (length < 1000 bp) ^17^.

### Bioinformatics analysis of long-read sequencing data

The long-read sequencing analysis was conducted similarly to the approaches used in our previous study^18^. Briefly, the clean long reads were mapped to a custom in-house reference genome, which includes the human genome (GRCh38) and the EBV type I genome (NCBI: NC_007605.1), using Minimap2 long-read mapper (v2.24)^19^. The resulting sequence alignment outputs in SAM format were then converted to BAM format and sorted by genomic coordinates using samtools (v1.9) ^20^ for subsequent SV identification and breakpoint detection. Additionally, to verify the existence of EBV sequences, the aligned reads were visualized in the Integrative Genomics Viewer (IGV)^21^ alongside the EBV genome. The SV caller SVIM (v 2.0.0)^22^ was employed to identify all types of SVs, including deletions, duplications, inversions, and translocations. Breakends (BND) associated with EBV were extracted, and the junction positions of human and EBV sequences were reported as the breakpoints for EBV integration. IGV was employed to load the alignment files and determine the annotations for corresponding integration events within GRCh38 and EBV reference genomes.

### PCR Verification and Sanger Sequencing of Integration Events

Primers were designed for specific integration events based on the Oxford Nanopore Technologies (ONT)’s long-read sequences upstream and downstream of the integration event, such that one primer binds on the integrated EBV sequence and the other binds to the human genome. Human GRch38/hg38 (GenBank ID: 883148) was used as the reference for the human genome. For NPC219, the integration event was amplified using primers 5’-CACAAATTATCTTCTGCCGTTTGAC-3’ and 5’-

CTCTCATGAACACTTATAGGCCATTAT-3’, expected product size of 786 bp. For C666-1, the integration event was amplified using primers 5’-AATGCCTCTCTTTGCCTACC-3’ and 5’-GCCAAATACCTTCTCCTTCCA-3, with an expected product size of 741 bp. All reactions were performed using 2X Taq FroggaMix (FroggaBio, Canada, Cat. FBTAQM), using the PCR conditions directed by the manufacturer, with an annealing temperature of 56°C for NPC219 and 52°C for C666-1. The PCR products were resolved on 2% agarose gels and the products were extracted using GenepHlow Gel/PCR Kit (Geneaid; SKU: DFH300). The purified products were then sequenced with Sanger sequencing platform at Genome Quebec using the 5’-CACAAATTATCTTCTGCCGTTTGAC-3’ and 5’-AATGCCTCTCTTTGCCTACC-3’ for NPC219 and C666-1, respectively.

### RNA-seq library construction

RNeasy® Mini kit from Qiagen was used to extract total RNA from snap frozen NPC219 tumor tissue. Library for whole transcriptomic sequencing was constructed by Genewiz (Azenta Life Science) and sequenced on Illumina platform.

### RNA sequencing data processing and analysis

RNA-sequencing bam files for the NPC cell lines were retrieved from a previous study^23^. Briefly, adapters and low-quality sequences were trimmed from the output reads. The trimmed short sequence reads were aligned to the hg38 reference genome using STAR aligner version 2.7.0 with the default parameters and quant mode for gene counts^24^. HT Seq was utilized for transcript quantification^25^. Then, DEGs were obtained using edgeR. An adjusted P-value of ≤0.05 was considered significant. To extract the DEGs, log2 fold changes greater than +2 and less than −2 were set as the cutoff values. To predict the interaction pattern of DEGs in EBV-positive samples, the PPI of DEGs was visualized using the Search Tool for the Retrieval of Interacting Genes/Proteins (STRING) database (https://string-db.org)^26^. Protein-protein interaction (PPI) networks constructed with a confidence score of ≥0.4 from the STRING databases were visualized by Cytoscape software (version 3.7.2)^27^. We removed the nodes with no interactions with other proteins. The connectivity degree of each node, which indicates the number of interactions of the corresponding gene, was examined by the CentiScaPe plugin in Cytoscape^28^. Nodes with a degree of connectivity ≥15 were labeled as hub genes. To identify fusion transcripts, we used the Arriba fusion finder to detect human-human and EBV-human fusions using the custom reference described earlier^29^. AnnoFuse was used to recognize potential oncogenic fusions involving only human transcripts^30^.

## Results

### Long-Read Sequencing of NPC for EBV Integration Analysis

Four EBV-positive samples were subjected to Nanopore long-read genome sequencing, including three cell lines and one tumor tissue (Table 1). After DNA extraction, the samples underwent quality testing, demonstrating satisfactory quantitative and qualitative results. The average DNA coverage across all samples was 10X (Supplementary Table S1). The median read length varied between approximately 2,280 bp and 10,300 bp across the four samples. On average, 47.1 Gb of sequences were generated per sample (Supplementary Table S1). The cleaned long reads were aligned to a custom in-house reference genome, consisting of the human genome and EBV type1 genomes, using the long-read mapper minimap2. Across the four samples, 3,056 EBV reads were detected, each longer than 1,000 bp. NPC219 displayed the greatest quantity of EBV reads, totaling 1,992 reads. The EBV read counts for the remaining samples ranged from 192 to 607 (Supplementary Table S1). The EBV content in each sample was visualized using IGV, demonstrating that reads from all samples spanned the entire length of the EBV genome (Figure 1A). Subsequently, SVIM was employed to detect structural variants. As a result, a total of 36 of integration breakpoints between EBV and human genomes were detected. We observed considerable diversity in integration sizes, spanning from 0.34 Kb to 112.1 Kb. To further validate the findings, we picked two representative breakpoints, one from the tumor and another from cell line sample for PCR amplification followed by sanger sequencing to verify the junction of EBV and human genome sequence. PCR confirmation revealed integration events for the two samples, confirming our long-read sequencing findings (Figure 1B-C). To further confirm the integration sequence, we sequenced representative PCR bands to reveal the EBV and human genome junctions (Figure 1B-C, chromatogram inlet). The sanger sequencing results matched with the long-read sequences at the breakpoints, showing both EBV and human base pairs.

**Figure 1:**
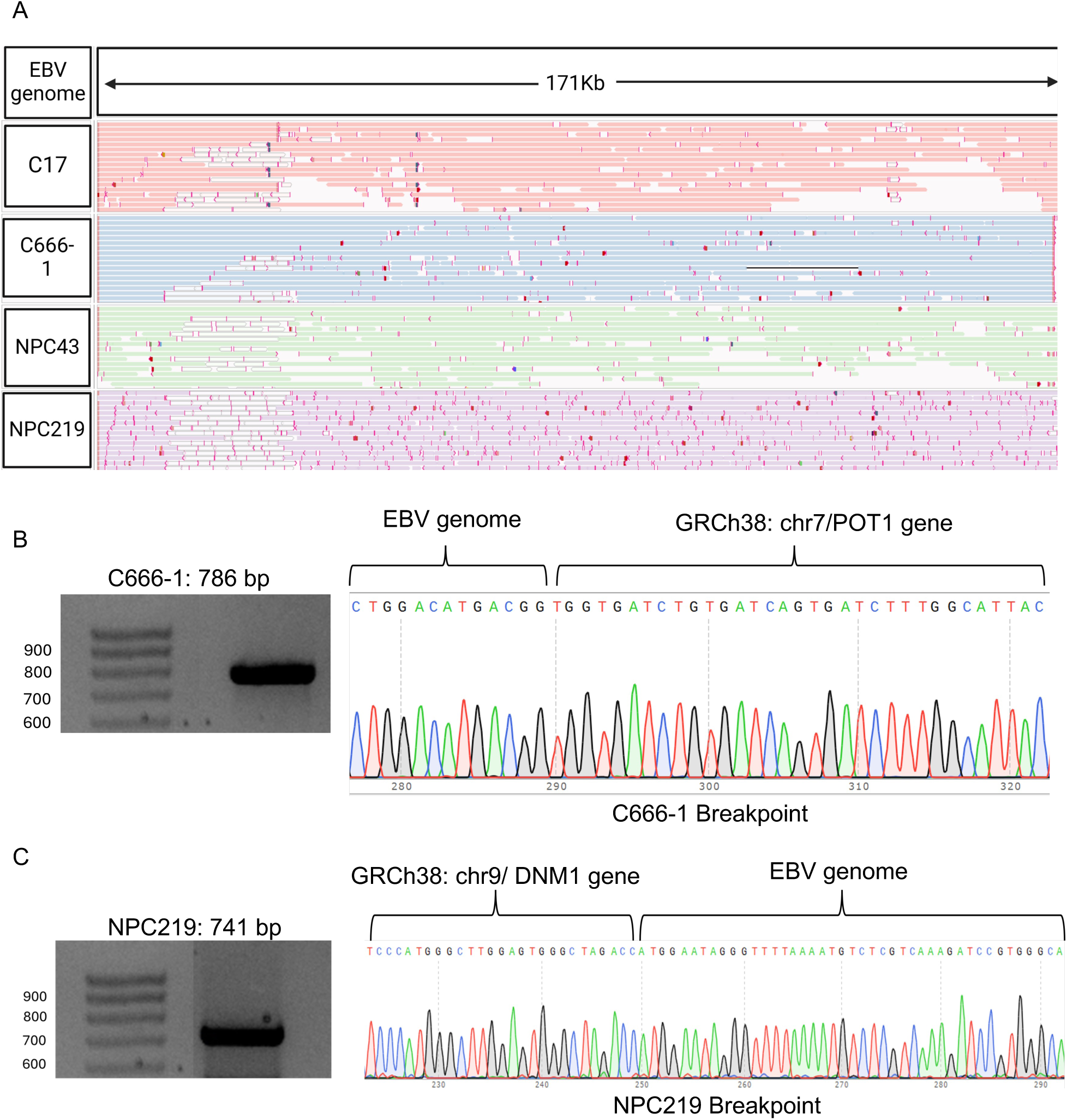
Long-Read Sequencing of NPC for EBV Integration Analysis. **a)** Representative ONT reads from C17, C666-1, NPC43, NPC219 samples aligned against the EBV genome: X-axis black arrows, the orientation of EBV genome from coordinates 1 to 171823 bps. The lines present the reads and are color-coded based on the samples. **b-c)** Representative PCR validation by amplification of two integration breakpoints for NPC219 (tumor) and C666-1 (cell line) samples with corresponding Sanger sequencing of PCR amplified integration events showing the junction of the EBV genome and the human genome.

**Table 1:**
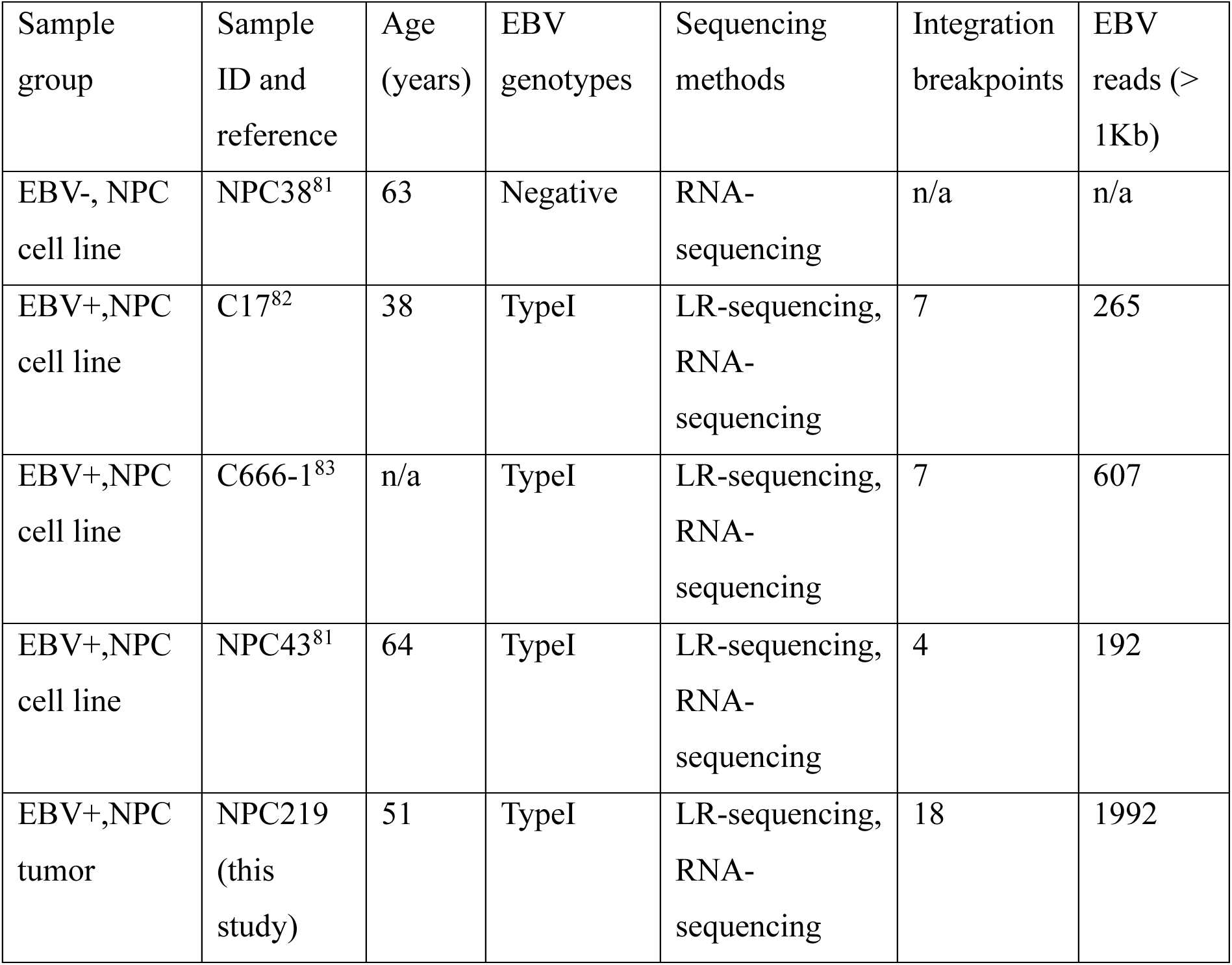
Clinical information of the Cell lines and tumor tissue.

### Breakpoints characterization and proximity of structural variants to integration breakpoints

The integration breakpoints were distributed across the human and virus genomes, with specific breakpoints concentrated in local hotspots (Figure 2A). Breakpoints were significantly enriched around EBNA1 and RPMS1 genes (Figure 2B). Previous findings have suggested that EBNA1 has the strongest negative correlation with viral RNA load, which is a signature for viral reactivation, aligning with its known role in inhibiting spontaneous viral reactivation ^31^. Similarly, RPMS1, a gene with virostatic properties (viral replication inhibitor), exhibits an inverse correlation with viral RNA load and has been shown to suppress lytic cycle induction ^32^ ^33^. Given the association between high EBV RNA load and cancer risk, potential inactivation of these genes could further raise the probability of cancer development ^11^. RPMS1 and EBNA1 potential virostatic functions might be silenced through integration-related disruptions, which can contribute to elevated EBV RNA loads and increased tumorigenesis. Furthermore, integration breakpoints within the same sample displayed patterns of occasional clustering. For instance, frequent breakpoints were localized around the genomic region involving PRP16 on chromosome 5 in sample C17, and there were clustering breakpoints around LINC00486 on chromosome 2 in sample Tumor_NPC219 (Figure 2A). EBV also showed a tendency to integrate near common fragile sites, with 36% of breakpoints found around repeat elements such as LINE, DNA transposons, and simple repeats (Figure 2C). Significantly, common fragile regions serve as preferred sites for EBV integration, acting as genomic hotspots for DNA damage and susceptible to genomic rearrangements^16,34,35^.

**Figure 2:**
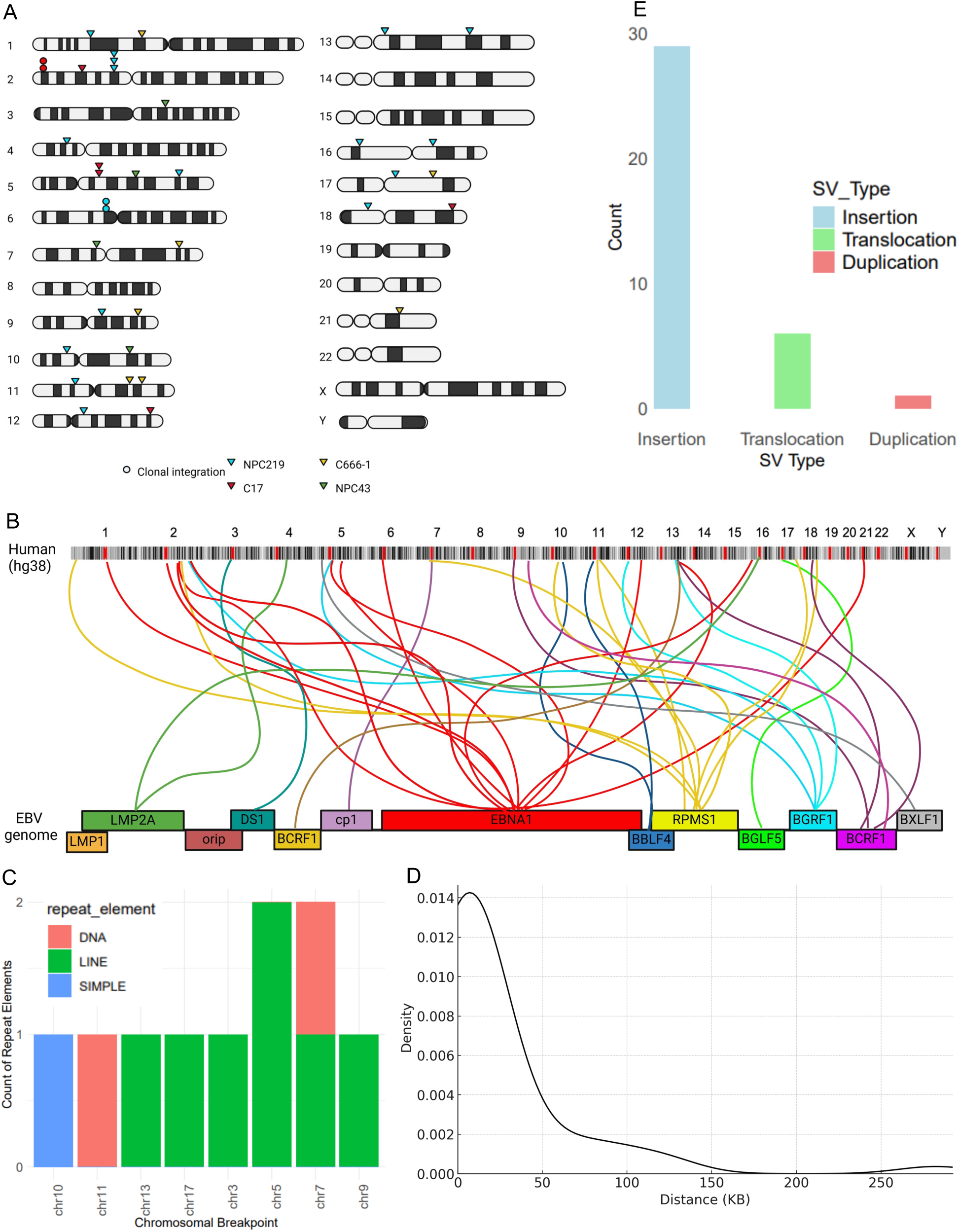
Breakpoints characterization and proximity of structural variants to integration breakpoints. **a)** Distribution of integration breakpoints across human and virus genomes. The colored link line indicates which human chromosome and virus gene the integration breakpoint had occurred on. Only EBV-related breakpoints are shown. **b)** Integration breakpoints distribution across the human genome with occasional clustering within samples. Triangles represent integration breakpoints, dots depict clonal integrations, and color keys indicate sample sources. **c)** Various types of repeat elements distributed across human breakpoints. **d)** Density plots of SVs distribution within 300Kb from upstream and downstream of integration breakpoint locations. **e)** The different types of SVs flanking the EBV integration breakpoints.

We investigated the co-localization of different SV types surrounding breakpoints to further analyze the genome instability induced by EBV integration. To control the genomic background, we mainly focused on genome windows spanning EBV integration sites where a window size of 500 Kb was selected. An elevated SV count was observed in a distance varying from 1 Kb to 30 Kb upstream and downstream of human-EBV breakpoints (Figure 2D). Among the five SV types, INS was specifically enriched near integration sites. We also witnessed translocation events colocalizing with breakpoints, further validating that EBV integration may increase the chromosomal translocation rate (Figure 2E). The findings indicate that EBV integration may cause genomic instability, resulting in an elevated incidence of structural variations with possible functional implications in the nasopharyngeal cancer progression.

### Types of EBV integration events

The occurrence of high-risk EBV DNA integration into the host genome is a critical molecular process affecting NPC development ^16^. We distinguished several forms of integrated EBV DNA by examining integration events on a case-by-case basis. These integrations mainly appeared in one of four forms: (I) single-fragment EBV integration, (II) integration of multiple EBV fragments from different genomic locations, (III) continuous long read containing majority of genome EBV (>80 Kb), and (IV) clonal EBV integrations. Due to the relatively long chimeric reads (2–120 Kb) obtained from the integration sites, we captured various integration modes. The most common type observed across samples was type I, characterized by the integration of a single fragment of the EBV genome followed by human DNA (Figure 3A). In samples C666-1, NPC219, and C17, we noted a higher prevalence of type II integrations, where multiple EBV segments from different locations had integrated (Figure 3B). Notably, in samples C17 and C666-1, we detected type III integration, where the majority of EBV genome integrations were detected (spanning 70–112 Kb), indicating that the entire EBV genome can integrate into the host DNA (Figure 3C). These samples also exhibited type I and type II integration events, supporting previous evidence that multiple EBV clones can coexist within a single tumor ^36^.

**Figure 3:**
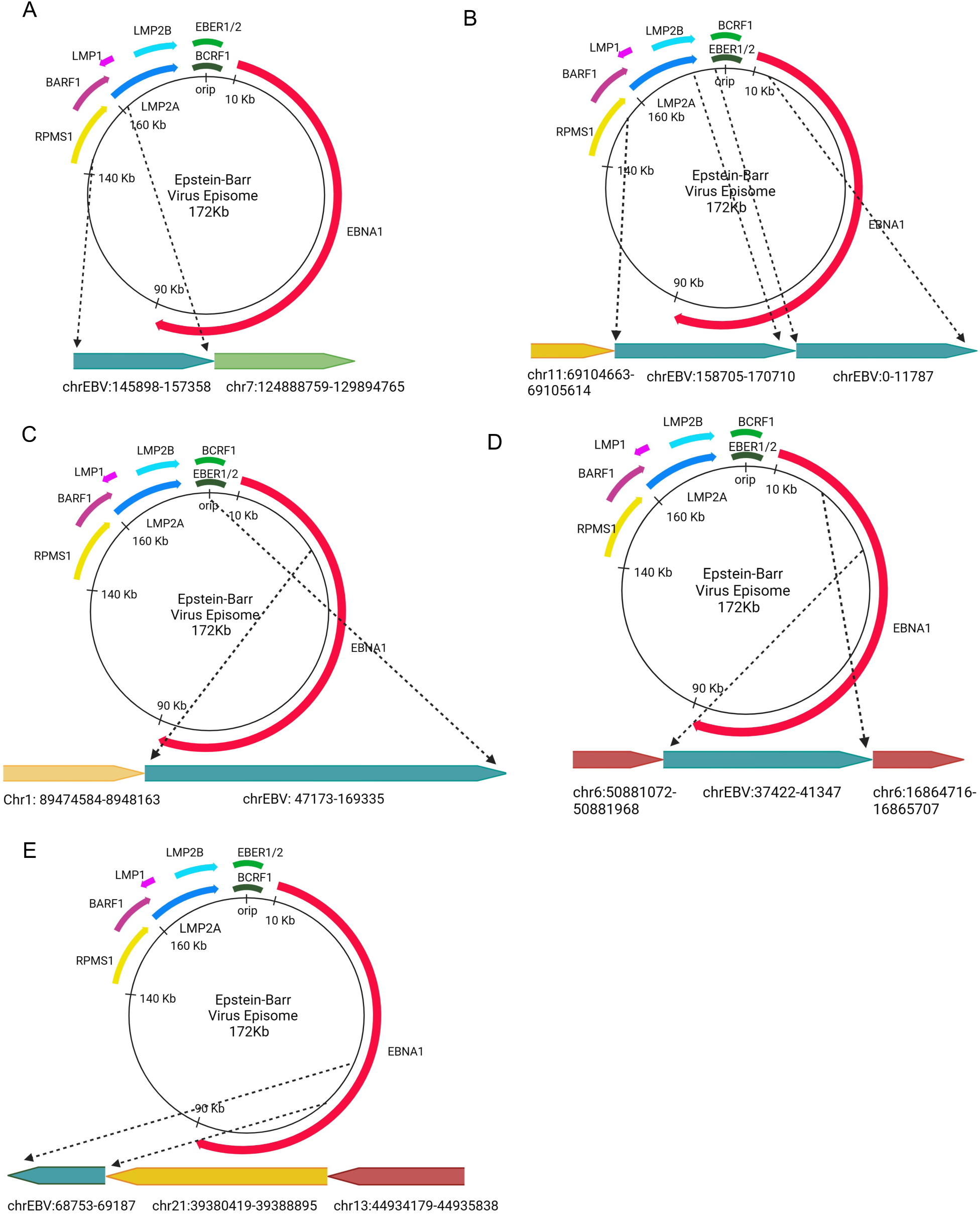
Types of EBV integration events and chromosomal translocation mediated by EBV DNA integration. **a)** Type (I) single-fragment EBV integration. **b)** Type (II) integration of multiple EBV fragments from different genomic locations. **c)** Type (III) integration of the majority of EBV genome (>80 Kb), an overflowing continuous segment containing the intact EBV genome. **d)** Type (IV) clonal EBV integration. **e)** Chromosomal translocation mediated by a segment of the EBV genome. In each panel, the two dashed arrows on the top indicate the EBV DNA fragment integrated into the human genome; the light blue and variously colored boxes on the bottom represent the EBV genome and human genome, respectively.

We also identified clonal integration events (type IV), defined by a series of chimeric reads in which the EBV DNA fragment was flanked by human genomic sequences. In NPC219 and C17, these clonal integrations involved EBV DNA flanked by human chromosomes 6 and 2, respectively (Figure 3D). The utilization of Nanopore technology, a sequencing method without amplification, facilitated direct, long-read sequencing of native DNA, enabling us to capture complex integration patterns. Each chimeric read likely corresponds to a single cell within the sequenced bulk cell population. The frequent observation of multiple types of integrated EBV DNA segments within individual tumors emphasizes the diversity and complexity of EBV integration in nasopharyngeal cancer.

### Chromosomal translocation mediated by EBV DNA integration

The analysis of long chimeric reads revealed that some EBV integration events were associated with inter-chromosomal and intra-chromosomal translocations, where viral DNA fused to host chromosomal segments. For example, in sample C666-1, a long chimeric read contained both a translocation of chromosomes 13 and 21 followed by a segment of EBV DNA (Figure 3E). Similarly, in sample NPC43, an EBV integration event coincided with a translocation between chromosomes 11 and 3 (Supplementary Table S2). A case of intra-chromosomal translocation was observed in sample NPC219, where EBV integration resulted in the fusion of two distinct regions of chromosome 6. Previous studies using different sequencing technologies have demonstrated similar viral-induced chromosomal translocations, particularly in cancers associated with human papillomavirus (HPV) ^37^. Short-read technologies often miss the complete scope of these events due to limited read length and the inability to capture the full spectrum of large chromosomal rearrangements, while long-read sequencing offers a more comprehensive view. The findings indicate that EBV integration not only affects genes around integration breakpoints but may also induce increased genomic instability involving chromosomal translocations.

### Gene expression alternations upon EBV integration

The integration of EBV potentially disrupted several genes, either directly at the integration sites or in close proximity to them. A total of 36 genes were annotated near integration breakpoints, consisting of 28 protein-coding genes and 8 non-coding RNAs (Supplementary Table S2). Integration hotspot regions were notably observed in the intergenic region downstream of the LINC00486 gene on chromosome 2 in sample NPC219 and upstream of PRR16 on chromosome 5 in sample C17 (Figure 2A). Other genes previously reported at known EBV integration hotspots included *ASAP2* and *TMEM132* ^38^. Long-read sequencing enabled precise mapping of both the breakpoints and the identification of the whole spectrum of the regions by EBV integration events. Across 36 integration events, the insertion of EBV into intergenic regions of human genome sequences was the most frequent phenomenon, with fragment lengths ranging from 340 to 112,163 bp. Additionally, integrated EBV DNA mediated inter-chromosomal and intra-chromosomal translocations, as observed in NPC219 and C666-1. Furthermore, large EBV segment integrations (type III) and translocation occurrences resulted in the disruption of multiple genes, totaling 41 genes identification around 36 breakpoints. Pathway enrichment analyses revealed that these genes were involved in crucial processes such as immunoregulatory interactions, cell cycle regulation, FGFR, and FoxO related signaling pathways (Figure 4A). To delve deeper into the functional consequences of EBV integration, these findings were combined with RNA-seq data. The expression levels of genes near integration breakpoints or within affected genomic regions were analyzed. By comparing the normalized expression counts of these genes across different samples, it was found that approximately 44% of the genes that contained EBV integration showed significant expression alternation upon EBV integration. Among the genes with altered expression were *CD96, GET1, POT1, DNM1, PIK3C2A, ABCD2, LRRC8B-D, ZMYM2, JAKMIP3*, and *ARHGAP27P*. Many of these genes have established associations with cancer, while others, such as *ARHGAP27P* and *JAKMIP3*, are relatively understudied and have not been previously linked to cancer.

**Figure 4:**
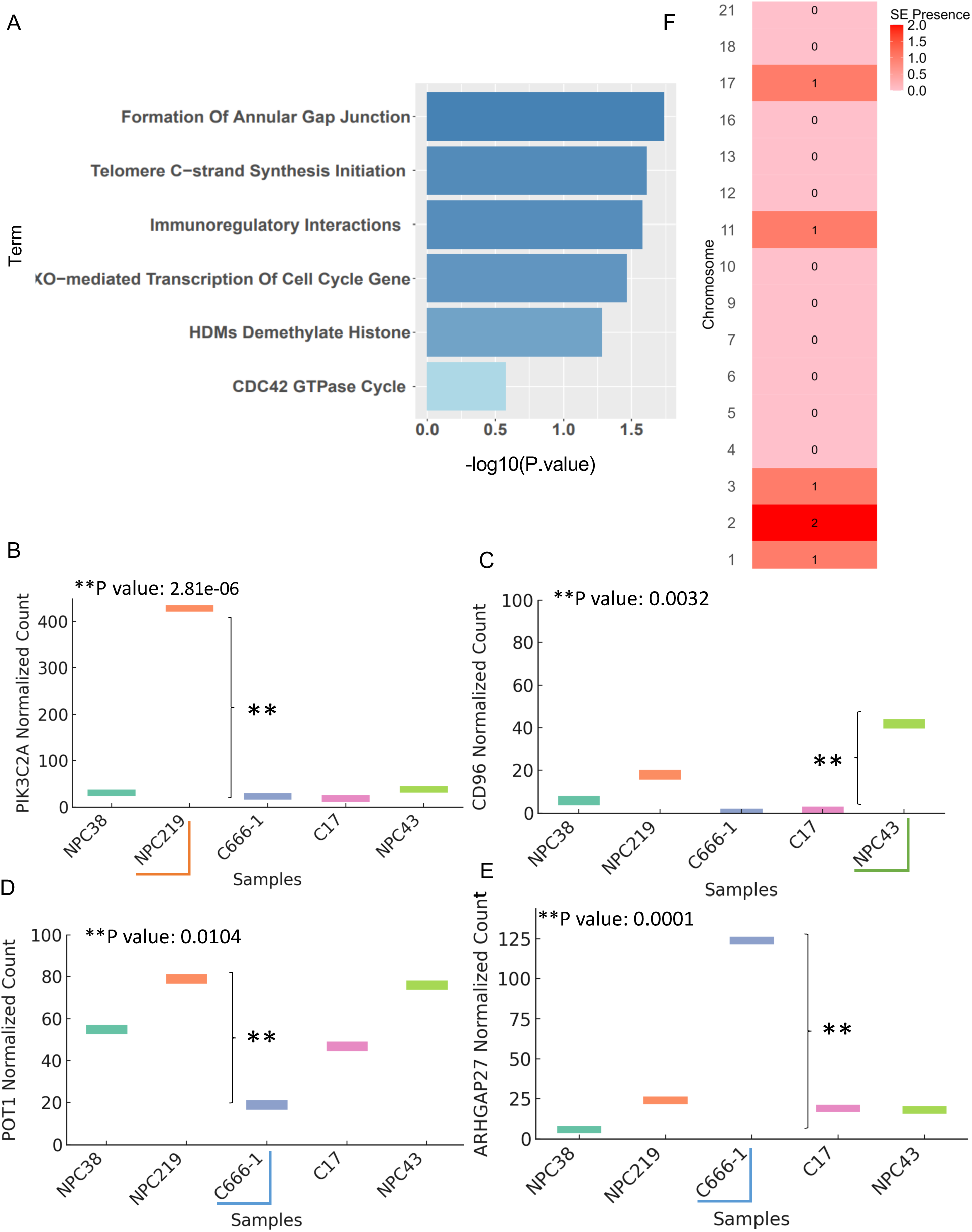
Gene expression alternations upon EBV integration. **a)** Bar graph of enriched pathways across 41 genes involved in 36 breakpoint events. The horizontal axis represented the P-value of enriched terms, and the vertical axis represented the enrichment of the functional pathways. **b-e)** Expression levels of four genes with EBV integration located within or in close proximity to them. The X-axis represents normalized count, and the Y-axis represents different samples. The sample with the gene integration is marked. One-sample t-test p-values for changes in gene expression were generated by evaluating the probability of the observed t-statistic under the assumption that the baseline expression follows a normal distribution. **f)** Heatmap indicating super-enhancers (SE) presence near integration breakpoints (distance < 100Kb) across human chromosomes.

We also detected a significant downregulation in the expression of several genes potentially disrupted by EBV integration, most notably tumor suppressor candidate such as *POT1*. Conversely, we identified instances where EBV integration was associated with the upregulation of genes with putative oncogenic functions. Among the most prominent of these are *CD96, ARHGAP27, POT1,* and *PIK3C2A* (Figure 4B-E, Supplementary Table S3). Upregulation of genes that are the target of EBV integration was also previously shown in a variety of EBV-positive cancer types, and it was associated with the super-enhancer (SE) region overlapping, suggesting that highly accessible chromatin may harbor more recurrent EBV integration events ^11^. In cancer cells, clusters of regulatory elements called SEs are generated near highly expressed genes important to tumorigenesis ^39^. We also detected instances of EBV integration sites significantly overlapped with SEs annotated in all the samples (Figure 4F). Moreover, the gene expression dysregulation, both upregulated and downregulated upon viral integration, has been shown with HPV viral integration^40^. This contrasting pattern of gene expression strongly indicates that EBV integration may facilitate tumorigenesis through a dual mechanism: downregulating tumor suppressor genes while simultaneously upregulating oncogenes.

### Differentially Expressed Genes (DEGs) analysis in EBV Positive and EBV Negative NPC Groups

High-throughput RNA sequencing was employed to investigate the impact of EBV on gene expression by comparing EBV-positive NPC samples to EBV-negative NPC sample NPC38. After normalizing raw counts, 275 differentially expressed genes (DEGs) were identified in the EBV-positive NPC samples compared to NPC38, of which 57 were upregulated and 218 were downregulated (Figure 5A and Supplementary Table S3). These results were obtained using a log2 fold change threshold of ≤ −2 or ≥ +2, along with an adjusted p-value of ≤ 0.05. We conducted Gene Ontology (GO) enrichment and canonical pathway analysis on the DEGs to gain further insights into the molecular landscape. Upregulated DEGs were significantly enriched in EBV-induced pathways, including CD4+ T cell activation, the JAK/STAT pathway, the NF-κB signaling pathway, and signaling cascades related to ribosomal and mitochondrial functions (Figure 5B). GO analysis also demonstrated a significant involvement of upregulated genes in oncogenic pathways in EBV-infected NPC samples, particularly through the p53 signaling cascade and pathways governing cell proliferation via cyclin-dependent factors. This finding underscores the role of EBV in modulating immune signaling, cell division and cellular metabolism further favoring NPC tumorigenesis. Conversely, the downregulated DEGs were enriched in pathways associated with epithelial-mesenchymal transition (EMT) through mechanisms involving focal adhesion, extracellular matrix (ECM) receptor interaction, and repression of gap junctions. This suggests that EBV may enhance the metastatic potential and invasiveness of tumors in NPC by promoting EMT. Furthermore, these downregulated genes were associated with immune evasion mechanisms, including suppression of the complement cascade and impaired chemotaxis (Figure 5C). Taken together, these results suggests that EBV could possibly regulate host-gene expression levels promoting oncogenesis in NPC.

**Figure 5:**
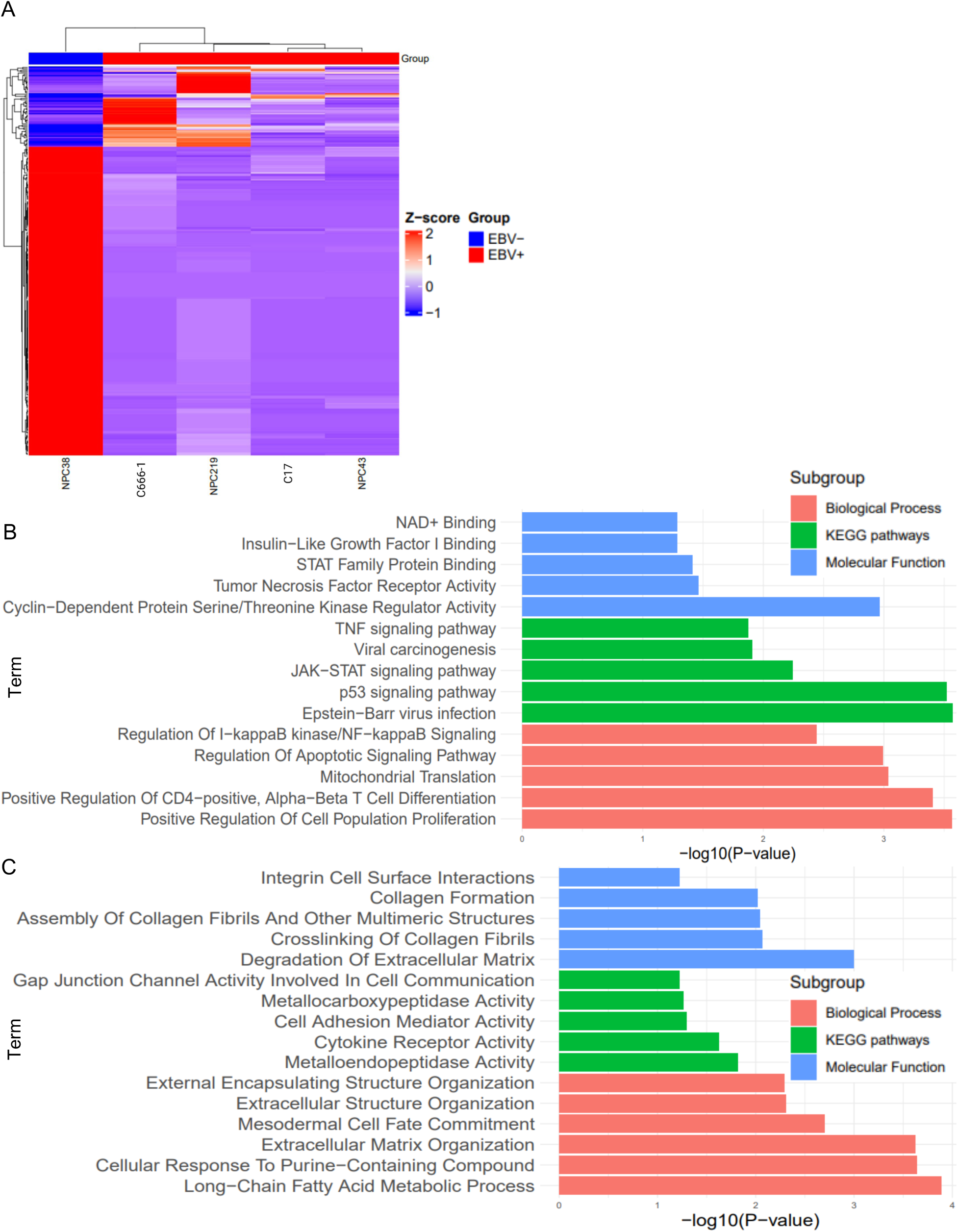
Differentially Expressed Genes (DEGs) analysis in EBV Positive and EBV Negative NPC Groups. **a)** Heatmap of two-way Hierarchical Clustering using the z-score computed for all the differentially expressed genes comparing EBV+ samples(red) versus EBV-control(blue). Each row represents a gene, and each column depicts a sample. Red indicates upregulation, and blue indicates downregulation. **b-c)** Bar graph of Gene Ontology (GO) enrichment analysis for upregulated (b) and downregulated (c) DEGs in three categories of pathways, molecular function, and biological process. The horizontal axis represented the P-value of enriched terms, and the vertical axis represented the enrichment of the functional pathways.

### Protein-Protein Interaction (PPI) Networks Analysis

To elucidate the downstream effects of host gene expression alternation that may follow EBV integration, we constructed two separate protein-protein interaction (PPI) networks, each offering a complementary viewpoint on this complex phenomenon. (1) a PPI network of DEGs derived from a comparison of EBV-positive and EBV-negative NPC samples, and (2) a mapping of interactions using the BioGRID interactome to elucidate direct interactions between EBV proteins and human DEGs, thereby uncovering potential mechanistic pathways of EBV-mediated influence on host cellular processes. The first group’s PPI network analysis revealed 196 nodes and 294 edges (Figure 6-Supplementary table S5). The second group network involving PPI interactions between EBV and human proteins, comprised 550 nodes and 680 edges (Figure 7-Supplementary table S6), demonstrating that only three of the total DEGs were direct interactors with EBV (CCND2, SNRPD2, FTSJ3) (Supplementary Table S6). Based on the degree of connectivity, we identified the top 15 genes for the first group as hub genes. Notably, BCL2 emerged as the key hub gene in the first group with a connectivity degree higher than 30 (Supplementary Table S5)

**Figure 6:**
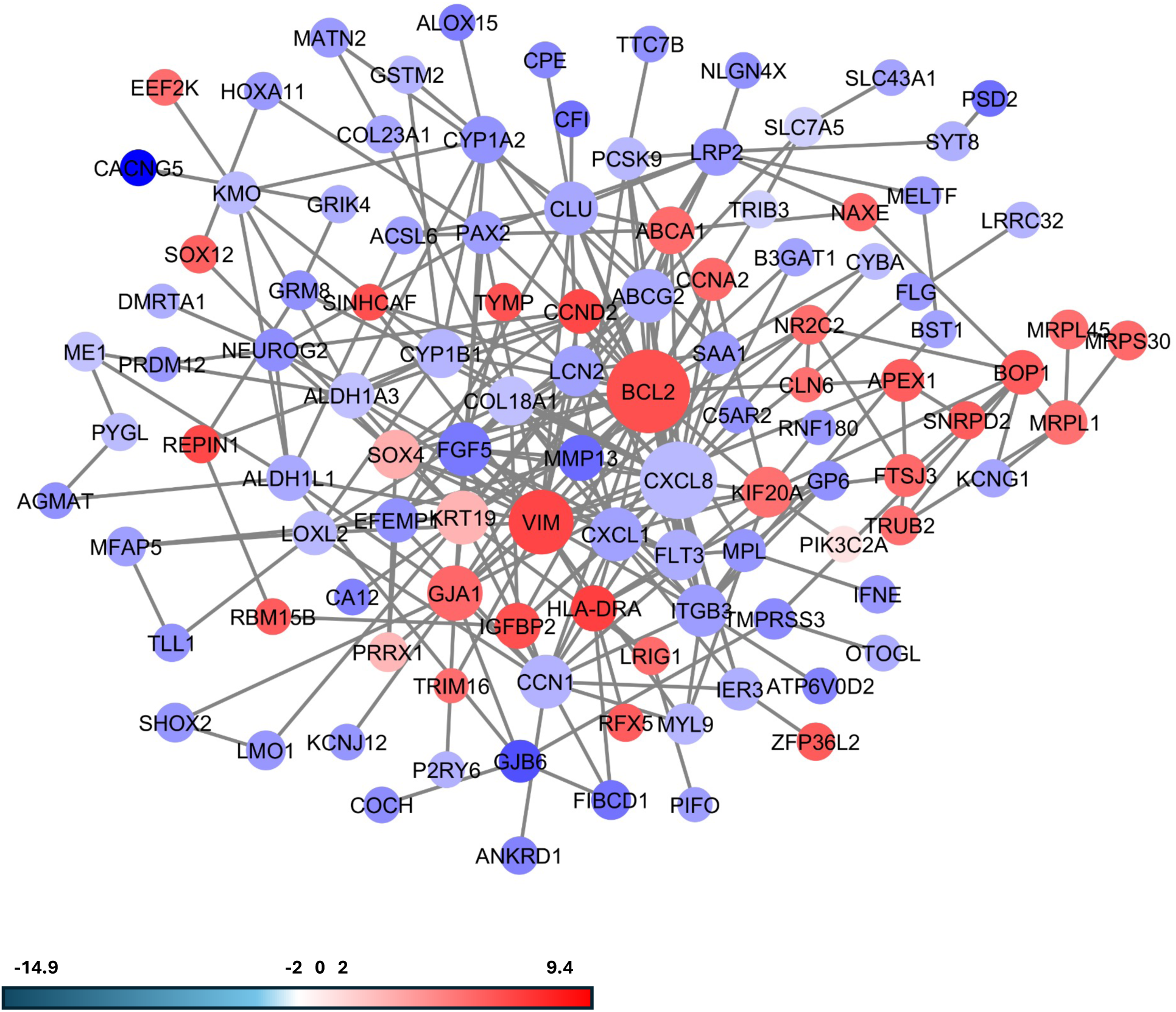
Protein-Protein Interaction (PPI) Networks Analysis. The PPI network was constructed with the DEGs. Circles/nodes present genes, and edges/lines indicate interactions. Disconnected nodes are hidden in the network. The size of each node represents the degree of connectivity based on the number of undirect and direct interactions used to identify the key hub genes. The red nodes are up-regulated genes, while the blue nodes represent the down-regulated genes.

**Figure 7:**
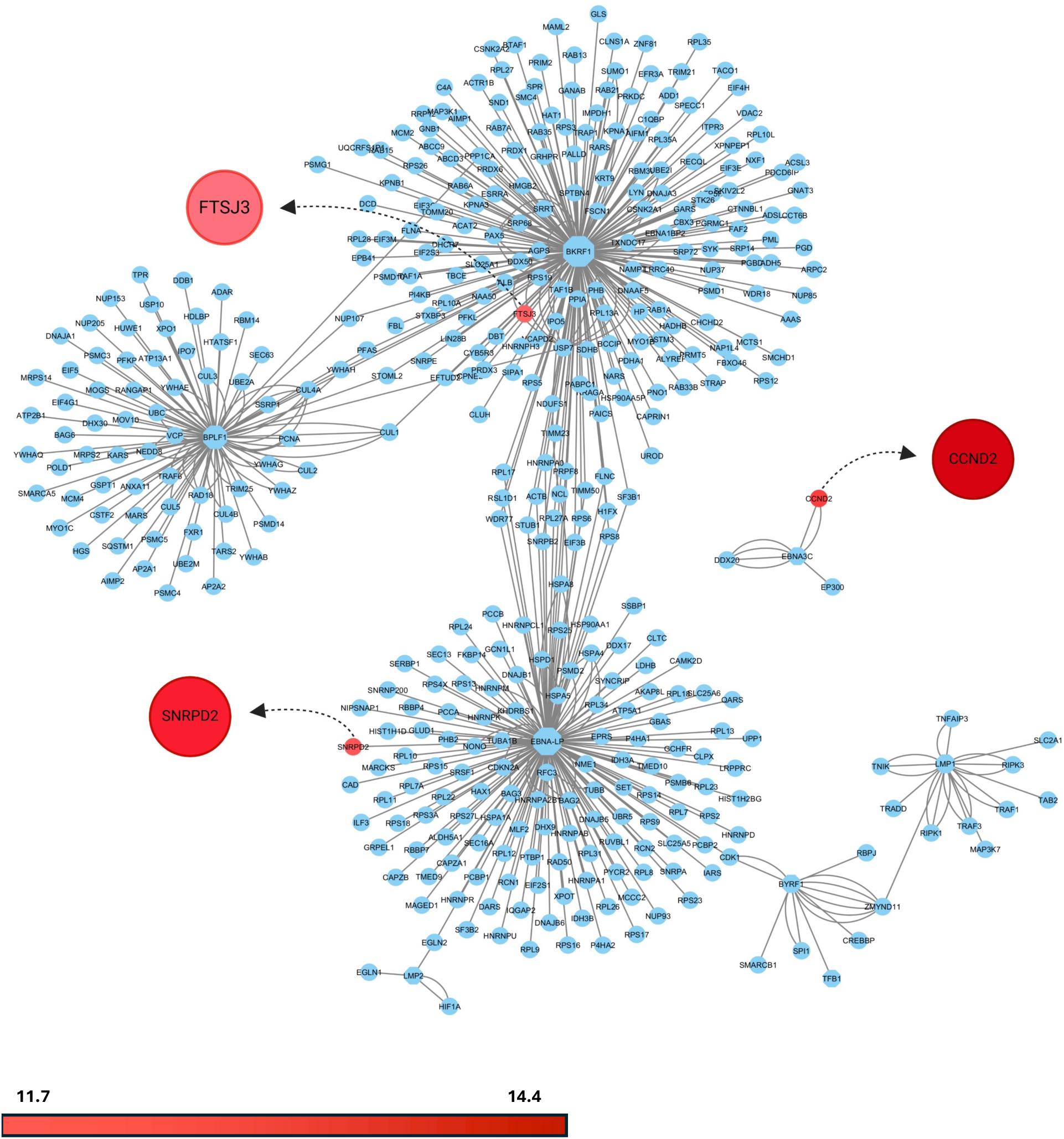
PPI network of interactome of main EBV proteins with human proteins. The nodes are colored based on the Log-FC value, the DEGs with direct EBV interaction are colored upregulated (red), and the nodes with light blue color are the interactors not included in the DEGs.

### EBV-human fusion transcripts existence in EBV-positive NPC

Gene fusions represent significant somatic alterations in cancer and are recognized as key drivers of tumorigenesis. After excluding gene fusions present in normal or para-cancerous tissues using annoFuse, we identified 319 unique fusion genes in NPC samples that were absent in the NPC38 sample (Figure 8A). Among these, 123 were intra-chromosome fusions, 196 were inter-chromosome fusions, and each chromosome had a distribution of fusion events. Among these, 123 were intra-chromosome fusions, 196 were inter-chromosome fusions, and each chromosome had a distribution of fusion events (Supplementary table S7). Seven fusion transcripts have been previously reported in The Cancer Genome Atlas (TCGA) Tumor Fusion Gene Database, including TEAD4-TULP3 in kidney renal clear cell carcinoma (KIRK) and NADK-GNB1 in lower-grade glioma (LGG), breast invasive carcinoma (BRCA), and head and neck squamous cell carcinoma (HNSCC) ^41^. We could also detect previously studied recurrent fusions in NPC such as UBR5–ZNF423 and RARS-MAD1L1 fusions that can facilitate tumorigenesis in sample C666-1 (Supplementary table S7) ^42,43^ .

**Figure 8:**
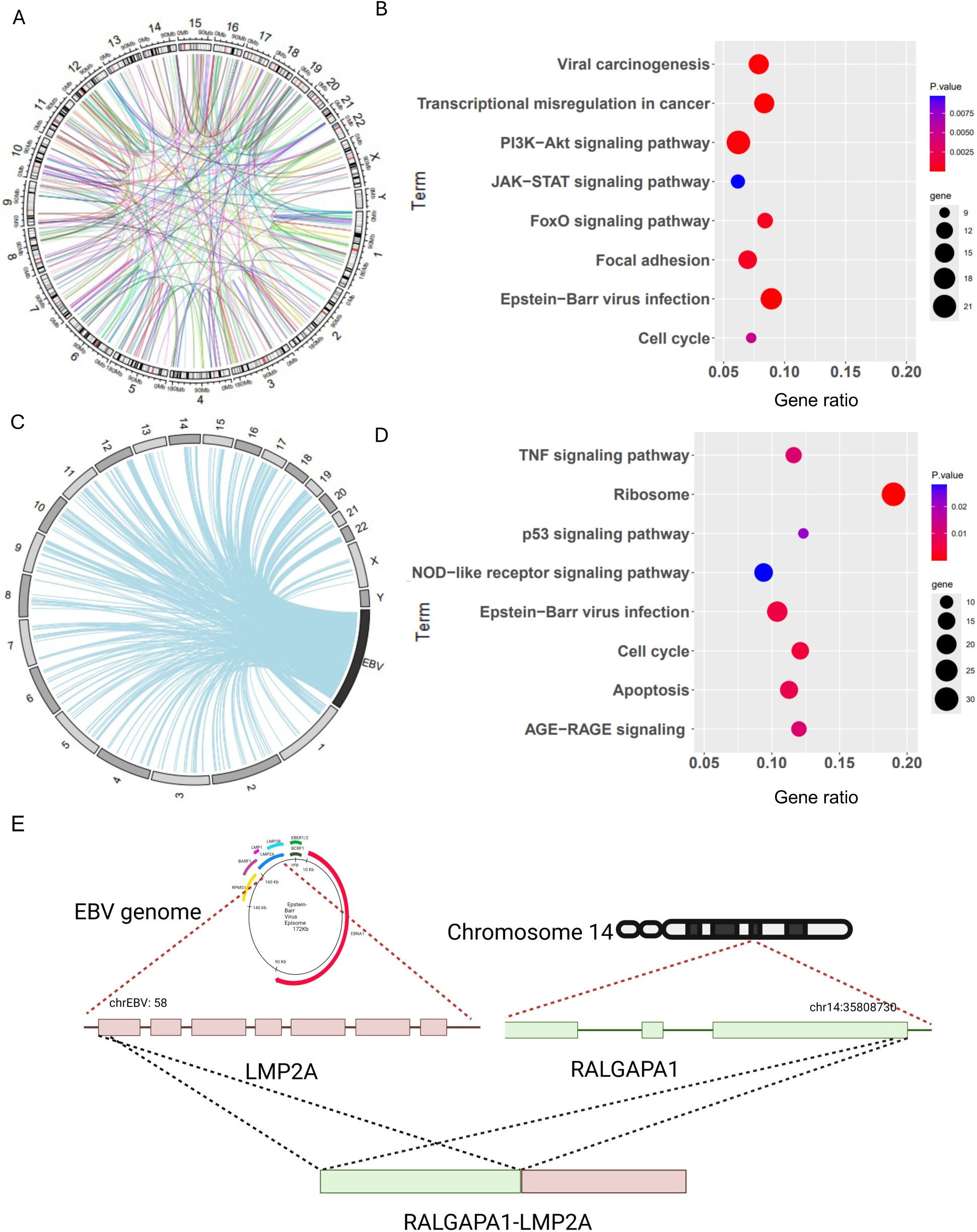
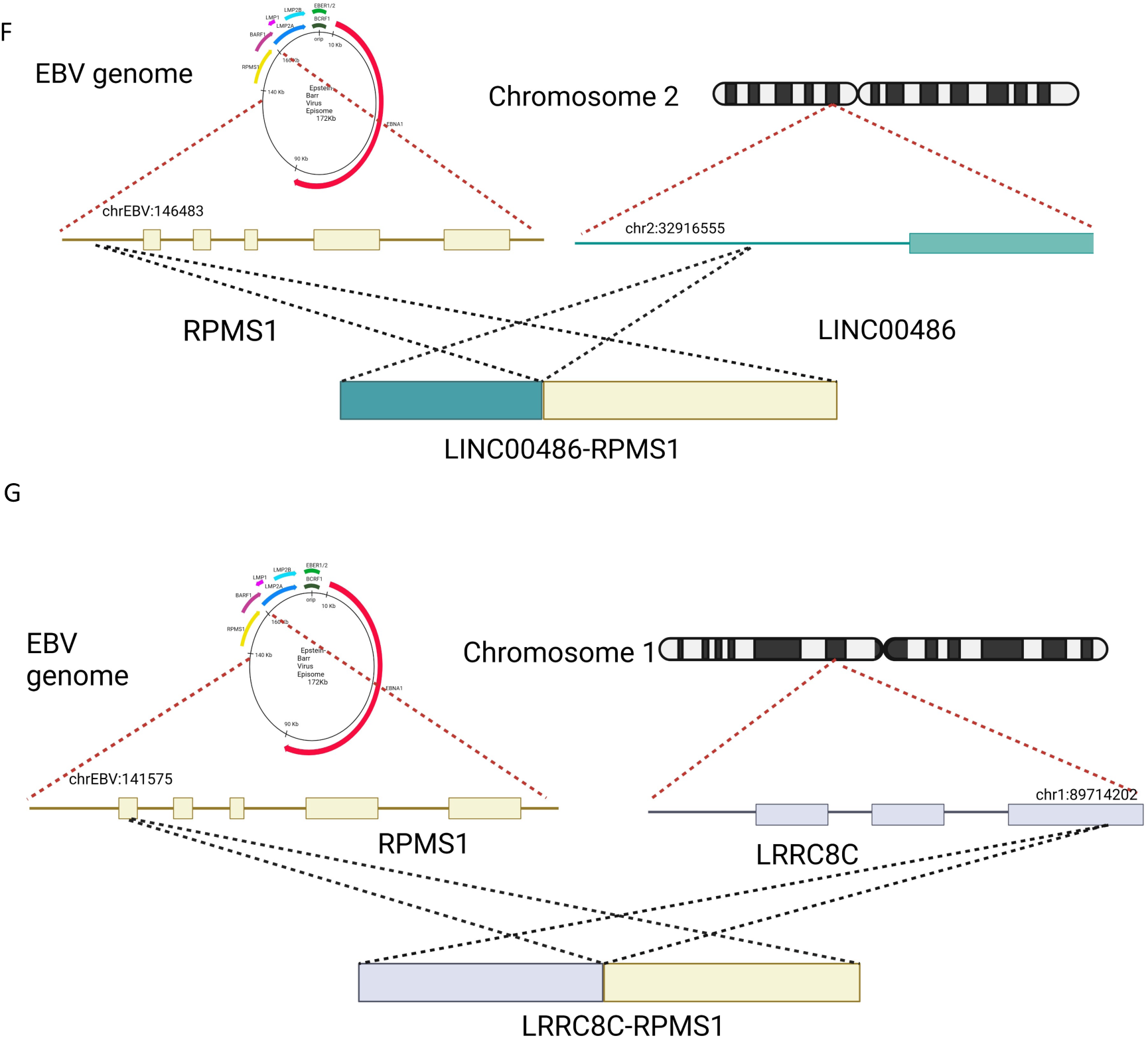
EBV-human fusion transcripts prevalent exist in EBV-related NPC. **a)** Circos plot showing the distribution of 320 fusion genes. The number in the outer ring was the human chromosome number. Each colored region of the middle ring represented the corresponding chromosome and its relative length. The curves between the human chromosomes represented the fusion gene connections. **b)** Bubble plots for enriched gene pathways for 519 genes involved in the 320 fusion events. The color of the nodes shown with a gradient from red to blue is according to the decreasing order of p value (p value < 0.05). The size of each node corresponds to several genes involved in each pathway. The horizontal axis represented the Gene Ratio (“Gene ratio” refers to the percentage of total candidate genes in the given pathway) of enriched terms, and the vertical axis represented the enrichment of the functional pathways. **c)** The Circos plot shows the integration of EBV genes (black bar) into human DNA (human chromosomes are shown in gray). Connecting lines between the human chromosomes and EBV represent the fusion gene events. **d)** Bubble plots for enriched gene pathways for 98 human genes involved in the 107 recurrent fusion events between sample NPC219 and C6661. The color of the nodes shown with a gradient from red to blue is according to the decreasing order of p value (p value < 0.05). The size of each node corresponds to the number of genes involved in each pathway. The horizontal axis represented the Gene Ratio of enriched terms, and the vertical axis represented the enrichment of the functional pathways. **e-g)** Schematic diagram of three fusion events formed by integration of EBV DNA into the human genome. The EBV genome and human chromosome associated with the fusion events were shown at the top. In each panel, the lines on the top indicate the EBV DNA fragment fused to the human genome. The structures of fusion transcripts of RALGAPA1-LMP2A (E), LINC00486-RPMS1(F), and LRRC8C-RPMS1(G) were shown below, with EBV fragments on the right and human fragments on the left side. The sizes of gene fragments were not in scale.

We performed an enrichment analysis of the fused genes to investigate the biological processes involved in the fusion events, which revealed that these genes were involved in EBV infection, B cell, and T cell receptor signaling pathways. There were also cancer related pathways including the cell cycle regulation, focal adhesion, transcriptional mis regulation in cancer, and Ras signaling pathway (Figure 8B). Among the four EBV+ NPC samples, we initially detected a novel EBV-human fusion RALGAPA1-LMP-2A in sample C666-1, which does not correspond to EBV integration detected in long read whole genome sequencing (Figure 8E). After we increased the mismatch tolerance of Arriba from 1% to 2% mismatches per read, new EBV-human fusion transcripts emerged in all the samples, with samples NPC43 and C17 with the lowest number of EBV fusions of 9 and 26 fusions, respectively. NPC219 and C666-1 displayed the highest number of fusions, 932 and 1171, respectively (Figure 8C and Supplementary Table S8). In this, two of them corresponded to the genes disrupted by EBV integration, including LRRC8C-RPMS1 for sample C666-1 and LINC00486-RPMS1 in sample NPC219 (Figure 8 F&G). A total of 105 fusions were recurrent between C666-1 and NPC219, and only DRD2-LMP1 fusion was recurrent in both C666-1 and NPC43. We also performed an enrichment analysis to investigate the biological processes involved in the recurrent fusions. The involved pathways included VEGF, P53, Notch, cell cycle, and EBV-related signaling pathways (Figure 8D).

## Discussion

In nasopharyngeal carcinogenesis, the integration of high-risk EBV DNA into the human genome is considered an essential molecular event ^34^. However, short-read NGS technologies have proven inadequate in providing a complete picture of the large-range EBV integration events and SVs accompanying EBV integration. To our knowledge, our study performed the first genome-wide analysis of EBV integration in four EBV-positive primary NPC tumors using Nanopore long-read sequencing and RNA sequencing technologies. Taking advantage of long-read sequencing, we could characterize at high resolution the various structures of the integrated EBV DNA segments.

We further revealed the relationship between EBV integration, the variety of SVs, and their potential consequences in NPC. Long-read sequencing analysis enabled us to decipher the full genomic length of EBV integrations in NPC for the first time. Our findings provide preliminary insights into the patterns of EBV integration into the human genome in NPC and its following impact on genomic instability and tumorigenesis. Characterization of integrated EBV DNA segments in the host genome will deepen our understanding of the pathogenic mechanism and the role of EBV in NPC. However, the present understanding of viral integration in EBV cancers is mostly based on Southern blot and short-read sequencing techniques ^44^ ^11^. These approaches provided a limited understanding of the whole virus-human integration processes in nasopharyngeal cancer. Using long-read sequencing methods, we attempted to fully define four unique kinds of integrated EBV DNA segments that are commonly present in integration events: simple integration, integration of multiple segments of EBV DNA, overflowing continuous segment containing the intact EBV genome, and clonal integrations. The existence of multiple EBV integration types in one sample supports the notion that EBV integration can coincide with episomal EBV genomes. This is best exemplified by the sample C666-1, as we detected several integration types such as type III with reads containing continuous segments of EBV and type II characterized with genome integration and several segments of EBV genome integrated into one locus.

We demonstrated that EBV integration is frequently associated with genomic instability, which may subsequently lead to an increased SV’s presence around integration sites, highlighting the EBV part in promoting chromosomal aberrations. Also, integration breakpoints in the same sample tended to be clustered in the host genome. Sample C17 was a good example where 28% of breakpoints were clustered on chromosome 5 (Figure 2B). A possible explanation for these intra-sample breakpoint clusters is that EBV integration induces local genomic instability, making this affected region more easily integrated by other EBV DNA fragments and facilitate structural variances formation. Nanopore long-read sequencing allowed us to capture complex integration patterns, including inter- and intra-chromosomal translocations involving EBV DNA. In these events, EBV either is placed as a linker in a clonal integration or is placed before or after an intra-chromosomal translocation. This is best illustrated in sample C666-1, where an integration event is related to a translocation involving chromosomes 13 and 21 (Figure 3E). Similarly, in sample NPC43, an integration event coincided with a translocation between chromosomes 11 and 3. These findings suggest that EBV integration influences genes around integration breakpoints and can induce genomic disruption through chromosomal translocation.

We identified 38 integration sites and 41 involved genes in 4 EBV-infected NPC samples. The integration breakpoints were distributed across both EBV and host genomes, breakpoints were frequently observed in regions such as EBNA1 and RPMS1, suggesting that disruption of these virostatic genes may positively regulate viral replication and cancer progression. Breakpoints were also co-localized near human genes such as *CD96, GET1, POT1, DNM1, PIK3C2A, ABCD2, LRRC8B-D, ZMYM2, ASAP2, LINC00486*, and *ARHGAP27P*, which are involved in various cellular pathways that contribute to tumorigenesis. Secondary genetic events in the host genome, resulting from EBV integration, are also essential for the development and progression of nasopharyngeal cancer. Combined with the corresponding RNA-seq data, we found that EBV integration may lead to host gene expression deregulation, altering key processes such as immune regulation, telomere maintenance, apoptosis, and cytoskeletal dynamics. CD96, for instance, is involved in immune checkpoint regulation ^45^, and its upregulated expression was reported in EBV-positive cancer, and it may contribute to immune evasion by EBV-positive tumor cells ^46^.Rho GTPase Activating Protein 27 (ARHGAP27) is a potential oncogene and acts as a Rho GTPase-activating protein^47^. It has been proposed that the ARHGAP27 gene could contribute to cancer development by disrupting the regulation of Rho/Rac/Cdc42-like GTPases^48^ ^49^ . According to the TIMER2.0 database, which estimates immune infiltration levels using TCGA data, there is a positive correlation between ARHGAP27 expression and the infiltration of CD4+ T cells in HNSCC samples, suggesting a potential role of ARHGAP27 in immune evasion ^50^ (Supplementary Figure S1). POT1 is a vital part of the shelterin complex and plays a crucial role in preserving telomere stability and securing genomic integrity. LMP1 has been shown to reversibly downregulate POT1 expression, resulting in progressive destabilization of shelterin three-dimensional configuration, which ultimately contributes to telomere dysfunction and the accumulation of intricate chromosomal rearrangements ^51^ ^52^ ^53^.

Neuroendocrine long-coiled-coil protein 2 (NECC2), also referred to as JAKMIP3 is a novel and understudied long-coiled-coil protein predominantly expressed in the central nervous system and endocrine tissues^54^. It was recently confirmed that NECC2 interacts with tyrosine kinase 2 (TYK2) and NECC2 silencing inhibited phosphorylation of Tyk2, and revealing NECC2-regulated Tyk2 activity ^55^. TYK2 is critical for the production of interferons and response to cytokines and plays a vital role in innate immunity and inflammation, including inflammatory conditions resulting from viral infections^56^. Interestingly, it is shown that LMP1 and BGLF2 interact with TYK2, inhibiting TYK2 phosphorylation and impairs type I IFN signaling to evade the host’s innate immune response^57,58^. Therefore, JAKMIP3 may act as a novel tumor suppressor in EBV+ NPCs through TYK2 activation. The involvement of known oncogenes such as CD96, ARHGAP27, and PIK3C2A, along with tumor suppressors like POT1 and JAKMIP3, highlights the dual mechanisms of oncogene activation and tumor suppressor inhibition that may modulate EBV tumorigenesis.

Protein-protein interaction network analysis offered a deeper insight into how the network’s central interactors may contribute to EBV-positive NPC tumorigenesis. B-cell lymphoma 2 (BCL2) emerged as the key hub gene in the first group of the PPI network, functioning as an essential apoptosis regulator^59^. Notably, BCL2 upregulation has been detected in EBV-positive NPCs^60,61^. BCL2 overexpression is induced via LMP1-A recruiting the TNF receptor (TNFR), leading to subsequential activation of three signaling cascades c-Jun N-terminal kinase (JNK), nuclear factor κB (NF-κB), and phosphoinositide 3-kinase (PI3K)/PKB/Akt, contributing to the enhancement of the pro-survival activity of the viral protein^37–39^. Kinesin family member 20A (KIF20A) is a crucial player in mitosis and cell division and was identified as a central interactor in the first group^65^. KIF20 lies downstream of VEGF signaling and mediates essential processes necessary for physiological and pathological vascular growth^66^. Importantly, enhanced expression of KIF20A was linked to worse overall survival (OS) and progression-free survival (PFS) among NPC patients^67^. Intriguingly, KIF20A is also a direct interactor of PIK3C2A, a member of PI3K class II, and is among the genes found to harbor EBV integration. This interaction likely occurs via the VEGF signaling pathway, where PIK3C2A can be involved in VEGF production stimulation through the PI3K/Akt signaling cascade^68^. PIK3C2A is a known oncogene that governs endosomal trafficking and cell migration through clathrin-mediated processes, which further corroborates its involvement in cancer development^69–71^.

Among the top hub genes directly interacting with EBV proteins, we identified FTSJ3, which interacts with EBNA1. FTSJ3, though understudied in cancer, has been shown to act as an RNA 2′-O-methyltransferase and has been associated with facilitating tumor immune evasion through epigenetic means, particularly by inhibiting abnormal double-stranded RNA-mediated type I interferon (IFN) responses^72, 73^. Furthermore, FTSJ3 has been demonstrated to support viral innate immune evasion via IFIH1/MDA5^74^. These discoveries imply that FTSJ3 could be a potential biomarker for EBV-positive NPC, considering its dual role in regulating the immune response and interacting with EBV proteins. In summary, these hub genes highlight the intricate nature of EBV interactions with host cellular mechanisms and genes, pinpointing specific targets through which EBV integration amplifies oncogenic signaling, enhances cell survival, and fosters tumor advancement. Comprehending these complex interactions can yield valuable insights for creating targeted therapies aimed at disrupting pivotal pathways that drive EBV-positive NPC.

Our data also revealed the presence of recurrent EBV-human fusion transcripts, further underscoring the complexity of EBV integration in NPC. The identification of novel fusion events, such as RALGAPA1-LMP-2A, LRRC8C-RPMS1, and LINC00486-RPMS1, highlights the diversity of integration outcomes and their potential impact on cellular functions. Notably, the RALGAPA1-LMP-2A fusion transcript did not correspond directly to the EBV integration breakpoints, which may be due to low coverage in the long-read genome sequencing or the possibility that fusion transcripts can form through read-through events during RNA transcription without requiring structural chromosomal changes^75^.

The RalGAPs are complexes composed of a catalytic RalGAPα1 or α2 subunit together with a common β subunit, and they regulate multiple cellular functions such as endocytosis, exocytosis, cell growth, and cytoskeletal reorganization via their effectors, including the exocyst complex ^76^. The RALGAPA1-LMP-2A fusion is particularly notable because the RalGAPα1 subunit typically exerts tumor suppressive effects through Ral GTPase inactivation ^77^. LMP-2A mimics constitutive B-cell receptor signaling, promoting cell survival and proliferation. When fused with RalGAPα1, LMP-2A may interfere with its tumor suppressive function, shifting its role towards oncogene. LRRC8C, a component of the LRRC8 ion channel receptor, plays a crucial role in the volume-regulated anion current (VRAC) in T cells, where it regulates T-cell function through the LRRC8C-STING-p53 signaling pathway, which acts as an inhibitory mechanism controlling immune responses^78,79^. RPMS1 fusion with LRRC8C may impair its virostatic function, increase EBV proliferation, and potentially contribute to immune evasion of EBV-positive cells.LINC00486, involved in the LINC00486-RPMS1 fusion, has been shown to inhibit proliferation and promote apoptosis in breast cancer by regulating miR-182-5p expression^80^. The fusion with RPMS1 may have oncogenic properties by disrupting the negative regulatory pathways involved, leading to uncontrolled cellular proliferation and apoptosis inhibition in NPC

Taken together, the discovery of recurrent EBV-human fusion transcripts in NPC samples highlights the intricate consequences of EBV integration on cellular function and tumorigenesis and adds a layer of which EBV can contribute to NPC tumorigenesis. These findings emphasize the role of fusion transcripts in shaping the oncogenic landscape of NPC and underscore the need for further investigation into their functional and clinical implications.

Several caveats of our study should be acknowledged. To begin with, our research faced challenges where some of the samples (particularly NPC tumor) have low coverage in long-read sequencing. Given the scarcity of NPC tumors, we performed this pilot study using well-established and characterized EBV-positive NPC cell lines, C666-1, NPC43 and C17, to determine the feasibility of this technology and our bioinformatics pipelines in uncovering EBV integration events. While we attempted to extract DNA and RNA from four NPC tumor samples, only one (NPC219) contained sufficient amount and quality for long-read sequencing and RNA-sequencing. Albeit its coverage is low, we were still able to detect integration events and validated the chimeric read in NPC219. While EBV-positive NPC cell lines (NPC43, C17, C666-1) are extensively employed as models for NPC research, studies on EBV integration breakpoints have been limited exclusively to the C666-1 cell line, with no investigations reported for NPC43 and C17. Further investigations, such as increased depth of sequencing and larger patient tumor sample sizes, are required to capture the full spectrum of EBV integration procedures and patterns in the human genome. Second, while we identified interesting genes and fusions due to EBV integration into the genome, the primary aim of this study was to shed more light on the overall characteristics of EBV genome integration, the various types of integration that may occur, and how this process might alter host gene expression. Consequently, additional investigations are required to confirm and explore the functions of the potential markers and fusions discussed in this study. Functional validation of some of these fusion genes and other host gene targets disrupted by EBV integration or interactions is required to shed more light on the NPC tumorigenesis.

In summary, our study provides a detailed characterization of EBV integration in nasopharyngeal carcinoma, highlighting the diverse integration patterns and their profound effects on host genomic stability and gene expression. The use of long-read sequencing technology has unveiled complex integration events that may contribute to tumor development and progression, offering new insights into the oncogenic potential of EBV in NPC. Future studies should focus on elucidating the precise mechanisms by which EBV integration influences host cellular pathways and exploring potential therapeutic strategies to mitigate the effects of viral integration in EBV-associated cancers.

## Author Contributions

ZK had full access to all of the data in the study and takes responsibility for the integrity of the data. *Concept and design*: ZK, AK, SKL. *Acquisition, analysis, or interpretation of data:* ZK, AK, AWYC, SCC and SKL. *Drafting of the manuscript*: ZK and SKL. *Critical revision of the manuscript for important intellectual content*: All authors. *Statistical analysis*: ZK. *Obtained funding*: SKL. *Administrative, technical, or material support*: AWYC, SCC and SKL. *Supervision*: SKL.

## Acknowledgements

The authors would like to thank Mr. Gary Levesque, Mr. Francois Lefebvre, Mr. Haig Djambazian, McGill Genome Centre for their help and support. The authors would also like to thank Dr. Swneke Donovan Bailey and Dr. Dae-Kyum Kim for their helpful insights and comments. We thank the late Prof. George Tsao for his generosity in providing the NPC cell lines (NPC43, C17, NPC38) used in this study.

## Data Availability

Available upon request and completing the necessary requirements.

---

**Supplementary Figure 1:**
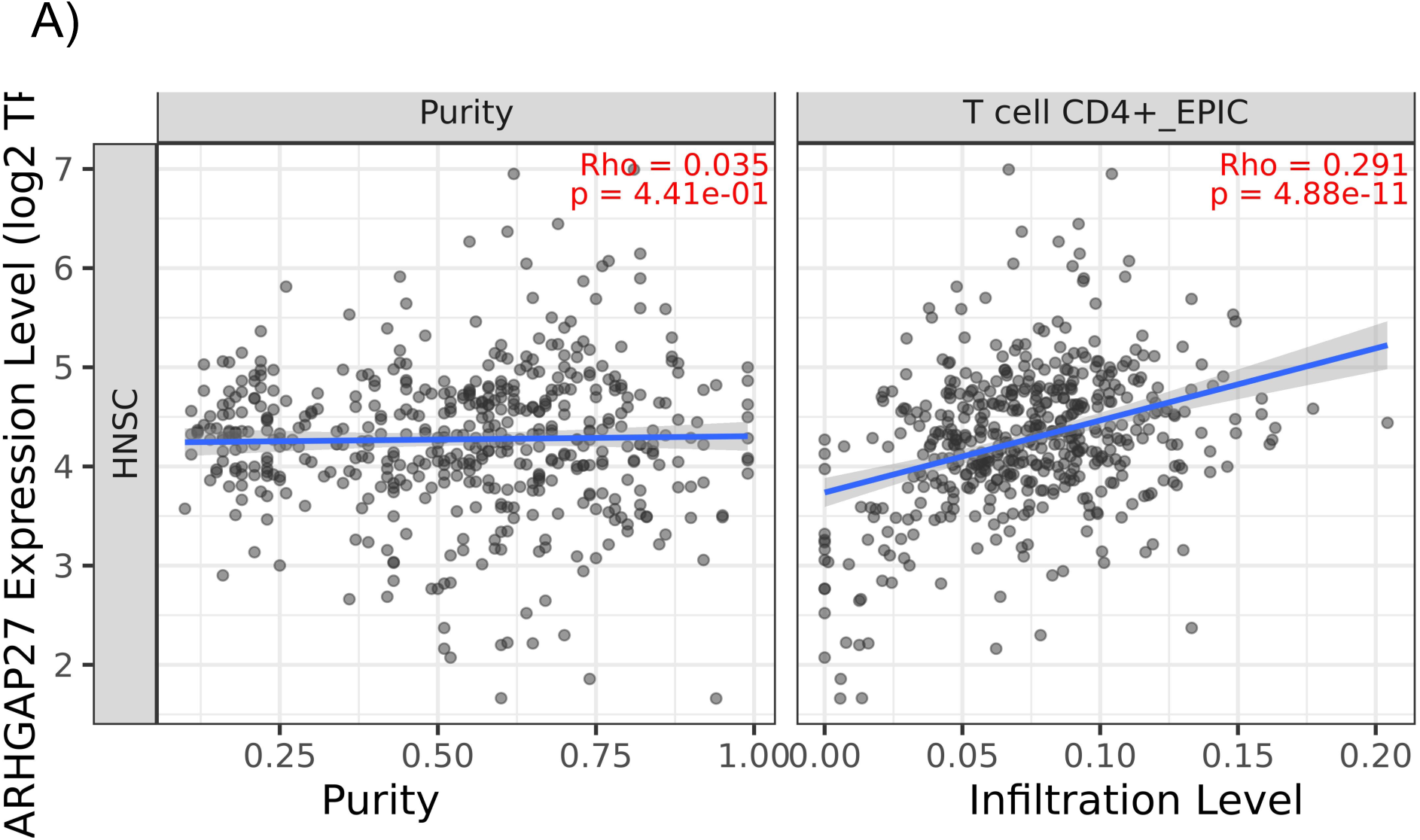
Correlation of *ARHGAP27* expression with CD4+ T cells immune infiltration in HNSCC from the TIMER2.0 database.

## References

(1) Sinha, S.; Winters, R.; Gajra, A. Nasopharyngeal Cancer. In StatPearls; StatPearls Publishing: Treasure Island (FL), 2024.

(2) Tang, L.-L.; Chen, W.-Q.; Xue, W.-Q.; He, Y.-Q.; Zheng, R.-S.; Zeng, Y.-X.; Jia, W.-H. Global Trends in Incidence and Mortality of Nasopharyngeal Carcinoma. Cancer Letters 2016, 374(1), 22–30. 10.1016/j.canlet.2016.01.040.

(3) Cancer Today. https://gco.iarc.who.int/today/ (accessed 2024-09-09).

(4) Blanchard, P.; Lee, A.; Marguet, S.; Leclercq, J.; Ng, W. T.; Ma, J.; Chan, A. T. C.; Huang, P.-Y.; Benhamou, E.; Zhu, G.; Chua, D. T. T.; Chen, Y.; Mai, H.-Q.; Kwong, D. L. W.; Cheah, S. L.; Moon, J.; Tung, Y.; Chi, K.-H.; Fountzilas, G.; Zhang, L.; Hui, E. P.; Lu, T.-X.; Bourhis, J.; Pignon, J. P. Chemotherapy and Radiotherapy in Nasopharyngeal Carcinoma: An Update of the MAC-NPC Meta-Analysis. The Lancet Oncology 2015, 16 (6), 645–655. 10.1016/S1470-2045(15)70126-9.

(5) Li, W.; Duan, X.; Chen, X.; Zhan, M.; Peng, H.; Meng, Y.; Li, X.; Li, X.-Y.; Pang, G.; Dou, X. Immunotherapeutic Approaches in EBV-Associated Nasopharyngeal Carcinoma. Front Immunol 2023, 13, 1079515. 10.3389/fimmu.2022.1079515.

(6) Lim, C. Y.; Ng, G. W. Y.; Goh, C. K.; Lee, M. K. C.; Cheong, I.; Ooi, E. E.; Liu, J.; West, R. B.; Loh, K. S.; Tay, J. K. Impact of High-Risk EBV Strains on Nasopharyngeal Carcinoma Gene Expression. Oral Oncology 2024, 157, 106941. 10.1016/j.oraloncology.2024.106941.

(7) Lung, M. L.; Cheung, A. K. L.; Ko, J. M. Y.; Lung, H. L.; Cheng, Y.; Dai, W. The Interplay of Host Genetic Factors and Epstein-Barr Virus in the Development of Nasopharyngeal Carcinoma. Chin J Cancer 2014, 33 (11), 556–568. 10.5732/cjc.014.10170.

(8) Okekpa, S. I.; Mydin, R. B. S. M. N.; Mangantig, E.; Azmi, N. S. A.; Zahari, S. N. S.; Kaur, G.; Musa, Y. Nasopharyngeal Carcinoma (NPC) Risk Factors: A Systematic Review and Meta-Analysis of the Association with Lifestyle, Diets, Socioeconomic and Sociodemographic in Asian Region. Asian Pac J Cancer Prev 2019, 20 (11), 3505–3514. 10.31557/APJCP.2019.20.11.3505.

(9) Zheng, H.; Dai, W.; Cheung, A. K. L.; Ko, J. M. Y.; Kan, R.; Wong, B. W. Y.; Leong, M. M. L.; Deng, M.; Kwok, T. C. T.; Chan, J. Y.-W.; Kwong, D. L.-W.; Lee, A. W.-M.; Ng, W. T.; Ngan, R. K. C.; Yau, C. C.; Tung, S.; Lee, V. H.-F.; Lam, K.-O.; Kwan, C. K.; Li, W. S.; Yau, S.; Chan, K.-W.; Lung, M. L. Whole-Exome Sequencing Identifies Multiple Loss-of-Function Mutations of NF-κB Pathway Regulators in Nasopharyngeal Carcinoma. Proc Natl Acad Sci U S A 2016, 113 (40), 11283– 11288. 10.1073/pnas.1607606113.

(10) Li, Y. Y.; Chung, G. T. Y.; Lui, V. W. Y.; To, K.-F.; Ma, B. B. Y.; Chow, C.; Woo, J. K. S.; Yip, K. Y.; Seo, J.; Hui, E. P.; Mak, M. K. F.; Rusan, M.; Chau, N. G.; Or, Y. Y. Y.; Law, M. H. N.; Law, P. P. Y.; Liu, Z. W. Y.; Ngan, H.-L.; Hau, P.-M.; Verhoeft, K. R.; Poon, P. H. Y.; Yoo, S.-K.; Shin, J.-Y.; Lee, S.-D.; Lun, S. W. M.; Jia, L.; Chan, A. W. H.; Chan, J. Y. K.; Lai, P. B. S.; Fung, C.-Y.; Hung, S.-T.; Wang, L.; Chang, A. M. V.; Chiosea, S. I.; Hedberg, M. L.; Tsao, S.-W.; van Hasselt, A. C.; Chan, A. T. C.; Grandis, J. R.; Hammerman, P. S.; Lo, K.-W. Exome and Genome Sequencing of Nasopharynx Cancer Identifies NF-κB Pathway Activating Mutations. Nat Commun 2017, 8, 14121. 10.1038/ncomms14121.

(11) Chakravorty, S.; Yan, B.; Wang, C.; Wang, L.; Quaid, J. T.; Lin, C. F.; Briggs, S. D.; Majumder, J.; Canaria, D. A.; Chauss, D.; Chopra, G.; Olson, M. R.; Zhao, B.; Afzali, B.; Kazemian, M. Integrated Pan-Cancer Map of EBV-Associated Neoplasms Reveals Functional Host–Virus Interactions. Cancer Research 2019, 79 (23), 6010–6023. https://doi.org/10.1158/0008-5472.CAN-19-0615.

(12) Perri, F.; Sabbatino, F.; Ottaiano, A.; Fusco, R.; Caraglia, M.; Cascella, M.; Longo, F.; Rega, R. A.; Salzano, G.; Pontone, M.; Marciano, M. L.; Piccirillo, A.; Montano, M.; Fasano, M.; Ciardiello, F.; Della Vittoria Scarpati, G.; Ionna, F. Impact of Epstein Barr Virus Infection on Treatment Opportunities in Patients with Nasopharyngeal Cancer. Cancers (Basel) 2023, 15 (5), 1626. 10.3390/cancers15051626.

(13) Chen, Y.-P.; Chan, A. T. C.; Le, Q.-T.; Blanchard, P.; Sun, Y.; Ma, J. Nasopharyngeal Carcinoma. The Lancet 2019, 394 (10192), 64–80. 10.1016/S0140-6736(19)30956-0.

(14) Weißbach, S.; Sys, S.; Hewel, C.; Todorov, H.; Schweiger, S.; Winter, J.; Pfenninger, M.; Torkamani, A.; Evans, D.; Burger, J.; Everschor-Sitte, K.; May-Simera, H. L.; Gerber, S. Reliability of Genomic Variants across Different Next-Generation Sequencing Platforms and Bioinformatic Processing Pipelines. BMC Genomics 2021, 22 (1), 62. 10.1186/s12864-020-07362-8.

(15) Janjetovic, S.; Hinke, J.; Balachandran, S.; Akyüz, N.; Behrmann, P.; Bokemeyer, C.; Dierlamm, J.; Murga Penas, E. M. Non-Random Pattern of Integration for Epstein-Barr Virus with Preference for Gene-Poor Genomic Chromosomal Regions into the Genome of Burkitt Lymphoma Cell Lines. Viruses 2022, 14 (1), 86. 10.3390/v14010086.

(16) Xu, M.; Zhang, W.-L.; Zhu, Q.; Zhang, S.; Yao, Y.; Xiang, T.; Feng, Q.-S.; Zhang, Z.; Peng, R.-J.; Jia, W.-H.; He, G.-P.; Feng, L.; Zeng, Z.-L.; Luo, B.; Xu, R.-H.; Zeng, M.-S.; Zhao, W.-L.; Chen, S.-J.; Zeng, Y.-X.; Jiao, Y. Genome-Wide Profiling of Epstein-Barr Virus Integration by Targeted Sequencing in Epstein-Barr Virus Associated Malignancies. Theranostics 2019, 9 (4), 1115–1124. 10.7150/thno.29622.

(17) Leger, A.; Leonardi, T. pycoQC, Interactive Quality Control for Oxford Nanopore Sequencing. Journal of Open Source Software 2019, 4 (34), 1236. 10.21105/joss.01236.

(18) Gauthier, M.-A.; Kadam, A.; Leveque, G.; Golabi, N.; Zeitouni, A.; Richardson, K.; Mascarella, M.; Sadeghi, N.; Loganathan, S. K. Long-Read Sequencing of Oropharyngeal Squamous Cell Carcinoma Tumors Reveal Diverse Patterns of High-Risk Human Papillomavirus Integration. Front. Oncol. 2023, 13. 10.3389/fonc.2023.1264646.

(19) Minimap2: pairwise alignment for nucleotide sequences | Bioinformatics | Oxford Academic. https://academic.oup.com/bioinformatics/article/34/18/3094/4994778 (accessed 2024-11-05).

(20) Li, H.; Handsaker, B.; Wysoker, A.; Fennell, T.; Ruan, J.; Homer, N.; Marth, G.; Abecasis, G.; Durbin, R.; Subgroup, 1000 Genome Project Data Processing. The Sequence Alignment/Map Format and SAMtools. Bioinformatics 2009, 25 (16), 2078. 10.1093/bioinformatics/btp352.

(21) Thorvaldsdóttir, H.; Robinson, J. T.; Mesirov, J. P. Integrative Genomics Viewer (IGV): High-Performance Genomics Data Visualization and Exploration. Briefings in Bioinformatics 2012, 14 (2), 178. 10.1093/bib/bbs017.

(22) SVIM: structural variant identification using mapped long reads | Bioinformatics | Oxford Academic. https://academic.oup.com/bioinformatics/article/35/17/2907/5298305 (accessed 2024-11-05).

(23) Chai, A. W. Y.; Yee, S. M.; Lee, H. M.; Abdul Aziz, N.; Yee, P. S.; Marzuki, M.; Wong, K. W.; Chiang, A. K. S.; Chow, L. K.-Y.; Dai, W.; Liu, T. F.; Tan, L. P.; Khoo, A. S. B.; Lo, K. W.; Lim, P. V. H.; Rajadurai, P.; Lightfoot, H.; Barthorpe, S.; Garnett, M. J.; Cheong, S. C. Establishment and Characterization of an Epstein-Barr Virus-Positive Cell Line from a Non-Keratinizing Differentiated Primary Nasopharyngeal Carcinoma. Cancer Res Commun 2024, 4 (3), 645–659. 10.1158/2767-9764.CRC-23-0341.

(24) Dobin, A.; Davis, C. A.; Schlesinger, F.; Drenkow, J.; Zaleski, C.; Jha, S.; Batut, P.; Chaisson, M.; Gingeras, T. R. STAR: Ultrafast Universal RNA-Seq Aligner. Bioinformatics 2012, 29 (1), 15. 10.1093/bioinformatics/bts635.

(25) Putri, G. H.; Anders, S.; Pyl, P. T.; Pimanda, J. E.; Zanini, F. Analysing High-Throughput Sequencing Data in Python with HTSeq 2.0. Bioinformatics 2022, 38 (10), 2943–2945. 10.1093/bioinformatics/btac166.

(26) Mering, C. von; Huynen, M.; Jaeggi, D.; Schmidt, S.; Bork, P.; Snel, B. STRING: A Database of Predicted Functional Associations between Proteins. Nucleic Acids Research 2003, 31 (1), 258. 10.1093/nar/gkg034.

(27) Shannon, P.; Markiel, A.; Ozier, O.; Baliga, N. S.; Wang, J. T.; Ramage, D.; Amin, N.; Schwikowski, B.; Ideker, T. Cytoscape: A Software Environment for Integrated Models of Biomolecular Interaction Networks. Genome Res 2003, 13 (11), 2498–2504. 10.1101/gr.1239303.

(28) Scardoni, G.; Tosadori, G.; Faizan, M.; Spoto, F.; Fabbri, F.; Laudanna, C. Biological Network Analysis with CentiScaPe: Centralities and Experimental Dataset Integration. F1000Research 2015, 3, 139. 10.12688/f1000research.4477.2.

(29) Uhrig, S.; Ellermann, J.; Walther, T.; Burkhardt, P.; Fröhlich, M.; Hutter, B.; Toprak, U. H.; Neumann, O.; Stenzinger, A.; Scholl, C.; Fröhling, S.; Brors, B. Accurate and Efficient Detection of Gene Fusions from RNA Sequencing Data. Genome Research 2021, 31 (3), 448. 10.1101/gr.257246.119.

(30) annoFuse: an R Package to annotate, prioritize, and interactively explore putative oncogenic RNA fusions | BMC Bioinformatics | Full Text. https://bmcbioinformatics.biomedcentral.com/articles/10.1186/s12859-020-03922-7 (accessed 2024-11-05).

(31) Sivachandran, N.; Wang, X.; Frappier, L. Functions of the Epstein-Barr Virus EBNA1 Protein in Viral Reactivation and Lytic Infection. J Virol 2012, 86 (11), 6146–6158. 10.1128/JVI.00013-12.

(32) Prince, S.; Keating, S.; Fielding, C.; Brennan, P.; Floettmann, E.; Rowe, M. Latent Membrane Protein 1 Inhibits Epstein-Barr Virus Lytic Cycle Induction and Progress via Different Mechanisms. J Virol 2003, 77 (8), 5000–5007. 10.1128/jvi.77.8.5000-5007.2003.

(33) Mp, R.; C, B.; M, W.; Ja, S.; D, N.; M, B. Silencing of Latent Membrane Protein 2B Reduces Susceptibility to Activation of Lytic Epstein-Barr Virus in Burkitt’s Lymphoma Akata Cells. The Journal of general virology 2007, 88 (Pt 5). 10.1099/vir.0.82790-0.

(34) Liao, X.; Zhu, W.; Zhou, J.; Li, H.; Xu, X.; Zhang, B.; Gao, X. Repetitive DNA Sequence Detection and Its Role in the Human Genome. Commun Biol 2023, 6 (1), 1–21. 10.1038/s42003-023-05322-y.

(35) Cao, Y.; Xie, L.; Shi, F.; Tang, M.; Li, Y.; Hu, J.; Zhao, L.; Zhao, L.; Yu, X.; Luo, X.; Liao, W.; Bode, A. M. Targeting the Signaling in Epstein–Barr Virus-Associated Diseases: Mechanism, Regulation, and Clinical Study. Sig Transduct Target Ther 2021, 6 (1), 1–33. 10.1038/s41392-020-00376-4.

(36) Peng, R.-J.; Han, B.-W.; Cai, Q.-Q.; Zuo, X.-Y.; Xia, T.; Chen, J.-R.; Feng, L.-N.; Lim, J. Q.; Chen, S.-W.; Zeng, M.-S.; Guo, Y.-M.; Li, B.; Xia, X.-J.; Xia, Y.; Laurensia, Y.; Chia, B. K. H.; Huang, H.-Q.; Young, K. H.; Lim, S. T.; Ong, C. K.; Zeng, Y.-X.; Bei, J.-X. Genomic and Transcriptomic Landscapes of Epstein-Barr Virus in Extranodal Natural Killer T-Cell Lymphoma. Leukemia 2019, 33 (6), 1451–1462. 10.1038/s41375-018-0324-5.

(37) Long-read sequencing unveils high-resolution HPV integration and its oncogenic progression in cervical cancer | Nature Communications. https://www.nature.com/articles/s41467-022-30190-1 (accessed 2024-10-07).

(38) Xiao, K.; Yu, Z.; Li, X.; Li, X.; Tang, K.; Tu, C.; Qi, P.; Liao, Q.; Chen, P.; Zeng, Z.; Li, G.; Xiong, W. Genome-Wide Analysis of Epstein-Barr Virus (EBV) Integration and Strain in C666-1 and Raji Cells. J Cancer 2016, 7 (2), 214–224. 10.7150/jca.13150.

(39) Hnisz, D.; Abraham, B. J.; Lee, T. I.; Lau, A.; Saint-André, V.; Sigova, A. A.; Hoke, H. A.; Young, R. A. Super-Enhancers in the Control of Cell Identity and Disease. Cell 2013, 155 (4), 934–947. 10.1016/j.cell.2013.09.053.

(40) Zhou, L.; Qiu, Q.; Zhou, Q.; Li, J.; Yu, M.; Li, K.; Xu, L.; Ke, X.; Xu, H.; Lu, B.; Wang, H.; Lu, W.; Liu, P.; Lu, Y. Long-Read Sequencing Unveils High-Resolution HPV Integration and Its Oncogenic Progression in Cervical Cancer. Nat Commun 2022, 13 (1), 2563. 10.1038/s41467-022-30190-1.

(41) X, H.; Q, W.; M, T.; F, B.; S, A.; K, Y.; Fm, L.; E, M.-L.; Sh, L.; S, Z.; Rgw, V. TumorFusions: An Integrative Resource for Cancer-Associated Transcript Fusions. Nucleic acids research 2018, 46 (D1). 10.1093/nar/gkx1018.

(42) Yang, L.; Zhu, J.-Y.; Zhang, J.-G.; Bao, B.-J.; Guan, C.-Q.; Yang, X.-J.; Liu, Y.-H.; Huang, Y.-J.; Ni, R.-Z.; Ji, L.-L. Far Upstream Element-Binding Protein 1 (FUBP1) Is a Potential c-Myc Regulator in Esophageal Squamous Cell Carcinoma (ESCC) and Its Expression Promotes ESCC Progression. Tumour Biol 2016, 37 (3), 4115–4126. 10.1007/s13277-015-4263-8.

(43) Chung, G. T.; Lung, R. W.; Hui, A. B.; Yip, K. Y.; Woo, J. K.; Chow, C.; Tong, C. Y.; Lee, S.-D.; Yuen, J. W.; Lun, S. W.; Tso, K. K.; Wong, N.; Tsao, S.-W.; Yip, T. T.; Busson, P.; Kim, H.; Seo, J.-S.; O’Sullivan, B.; Liu, F.-F.; To, K.-F.; Lo, K.-W. Identification of a Recurrent Transforming UBR5– ZNF423 Fusion Gene in EBV-Associated Nasopharyngeal Carcinoma. J Pathol 2013, 231 (2), 158–167. 10.1002/path.4240.

(44) Ohshima, K.; Suzumiya, J.; Kanda, M.; Kato, A.; Kikuchi, M. Integrated and Episomal Forms of Epstein–Barr Virus (EBV) in EBV Associated Disease. Cancer Letters 1998, 122 (1), 43–50. 10.1016/S0304-3835(97)00368-6.

(45) Thorsson, V.; Gibbs, D. L.; Brown, S. D.; Wolf, D.; Bortone, D. S.; Ou Yang, T.-H.; Porta-Pardo, E.; Gao, G. F.; Plaisier, C. L.; Eddy, J. A.; Ziv, E.; Culhane, A. C.; Paull, E. O.; Sivakumar, I. K. A.; Gentles, A. J.; Malhotra, R.; Farshidfar, F.; Colaprico, A.; Parker, J. S.; Mose, L. E.; Vo, N. S.; Liu, J.; Liu, Y.; Rader, J.; Dhankani, V.; Reynolds, S. M.; Bowlby, R.; Califano, A.; Cherniack, A. D.; Anastassiou, D.; Bedognetti, D.; Mokrab, Y.; Newman, A. M.; Rao, A.; Chen, K.; Krasnitz, A.; Hu, H.; Malta, T. M.; Noushmehr, H.; Pedamallu, C. S.; Bullman, S.; Ojesina, A. I.; Lamb, A.; Zhou, W.; Shen, H.; Choueiri, T. K.; Weinstein, J. N.; Guinney, J.; Saltz, J.; Holt, R. A.; Rabkin, C. S.; Cancer Genome Atlas Research Network; Lazar, A. J.; Serody, J. S.; Demicco, E. G.; Disis, M. L.; Vincent, B. G.; Shmulevich, I. The Immune Landscape of Cancer. Immunity 2018, 48 (4), 812–830.e14. 10.1016/j.immuni.2018.03.023.

(46) Salnikov, M.; Prusinkiewicz, M. A.; Lin, S.; Ghasemi, F.; Cecchini, M. J.; Mymryk, J. S. Tumor-Infiltrating T Cells in EBV-Associated Gastric Carcinomas Exhibit High Levels of Multiple Markers of Activation, Effector Gene Expression, and Exhaustion. Viruses 2023, 15 (1), 176. 10.3390/v15010176.

(47) Katoh, Y.; Katoh, M. Identification and Characterization of ARHGAP27 Gene in Silico. Int J Mol Med 2004, 14 (5), 943–947.

(48) Permuth-Wey, J.; Lawrenson, K.; Shen, H. C.; Velkova, A.; Tyrer, J. P.; Chen, Z.; Lin, H.-Y.; Chen, Y. A.; Tsai, Y.-Y.; Qu, X.; Ramus, S. J.; Karevan, R.; Lee, J.; Lee, N.; Larson, M. C.; Aben, K. K.; Anton-Culver, H.; Antonenkova, N.; Antoniou, A. C.; Armasu, S. M. Australian Cancer Study Australian Ovarian Cancer Study; Bacot, F.; Baglietto, L.; Bandera, E. V.; Barnholtz-Sloan, J.; Beckmann, M. W.; Birrer, M. J.; Bloom, G.; Bogdanova, N.; Brinton, L. A.; Brooks-Wilson, A.; Brown, R.; Butzow, R.; Cai, Q.; Campbell, I.; Chang-Claude, J.; Chanock, S.; Chenevix-Trench, G.; Cheng, J. Q.; Cicek, M. S.; Coetzee, G. A.; Consortium of Investigators of Modifiers of BRCA1/2; Cook, L. S.; Couch, F. J.; Cramer, D. W.; Cunningham, J. M.; Dansonka-Mieszkowska, A.; Despierre, E.; Doherty, J. A.; Dörk, T.; du Bois, A.; Dürst, M.; Easton, D. F.; Eccles, D.; Edwards, R.; Ekici, A. B.; Fasching, P. A.; Fenstermacher, D. A.; Flanagan, J. M.; Garcia-Closas, M.; Gentry-Maharaj, A.; Giles, G. G.; Glasspool, R. M.; Gonzalez-Bosquet, J.; Goodman, M. T.; Gore, M.; Górski, B.; Gronwald, J.; Hall, P.; Halle, M. K.; Harter, P.; Heitz, F.; Hillemanns, P.; Hoatlin, M.; Høgdall, C. K.; Høgdall, E.; Hosono, S.; Jakubowska, A.; Jensen, A.; Jim, H.; Kalli, K. R.; Karlan, B. Y.; Kaye, S. B.; Kelemen, L. E.; Kiemeney, L. A.; Kikkawa, F.; Konecny, G. E.; Krakstad, C.; Kjaer, S. K.; Kupryjanczyk, J.; Lambrechts, D.; Lambrechts, S.; Lancaster, J. M.; Le, N. D.; Leminen, A.; Levine, D. A.; Liang, D.; Lim, B. K.; Lin, J.; Lissowska, J.; Lu, K. H.; Lubiński, J.; Lurie, G.; Massuger, L. F. A. G.; Matsuo, K.; McGuire, V.; McLaughlin, J. R.; Menon, U.; Modugno, F.; Moysich, K. B.; Nakanishi, T.; Narod, S. A.; Nedergaard, L.; Ness, R. B.; Nevanlinna, H.; Nickels, S.; Noushmehr, H.; Odunsi, K.; Olson, S. H.; Orlow, I.; Paul, J.; Pearce, C. L.; Pejovic, T.; Pelttari, L. M.; Pike, M. C.; Poole, E. M.; Raska, P.; Renner, S. P.; Risch, H. A.; Rodriguez-Rodriguez, L.; Rossing, M. A.; Rudolph, A.; Runnebaum, I. B.; Rzepecka, I. K.; Salvesen, H. B.; Schwaab, I.; Severi, G.; Shridhar, V.; Shu, X.-O.; Shvetsov, Y. B.; Sieh, W.; Song, H.; Southey, M. C.; Spiewankiewicz, B.; Stram, D.; Sutphen, R.; Teo, S.-H.; Terry, K. L.; Tessier, D. C.; Thompson, P. J.; Tworoger, S. S.; van Altena, A. M.; Vergote, I.; Vierkant, R. A.; Vincent, D.; Vitonis, A. F.; Wang-Gohrke, S.; Palmieri Weber, R.; Wentzensen, N.; Whittemore, A. S.; Wik, E.; Wilkens, L. R.; Winterhoff, B.; Woo, Y. L.; Wu, A. H.; Xiang, Y.-B.; Yang, H. P.; Zheng, W.; Ziogas, A.; Zulkifli, F.; Phelan, C. M.; Iversen, E.; Schildkraut, J. M.; Berchuck, A.; Fridley, B. L.; Goode, E. L.; Pharoah, P. D. P.; Monteiro, A. N. A.; Sellers, T. A.; Gayther, S. A. Identification and Molecular Characterization of a New Ovarian Cancer Susceptibility Locus at 17q21.31. Nat Commun 2013, 4, 1627. 10.1038/ncomms2613.

(49) Yang, Y.; Wu, L.; Shu, X.; Lu, Y.; Shu, X.-O.; Cai, Q.; Beeghly-Fadiel, A.; Li, B.; Ye, F.; Berchuck, A.; Anton-Culver, H.; Banerjee, S.; Benitez, J.; Bjørge, L.; Brenton, J. D.; Butzow, R.; Campbell, I. G.; Chang-Claude, J.; Chen, K.; Cook, L. S.; Cramer, D. W.; DeFazio, A.; Dennis, J.; Doherty, J. A. ; Dörk, T.; Eccles, D. M.; Edwards, D. V.; Fasching, P. A.; Fortner, R. T.; Gayther, S. A.; Giles, G. G.; Glasspool, R. M.; Goode, E. L.; Goodman, M. T.; Gronwald, J.; Harris, H. R.; Heitz, F.; Hildebrandt, M. A.; Høgdall, E.; Høgdall, C. K.; Huntsman, D. G.; Kar, S. P.; Karlan, B. Y.; Kelemen, L. E.; Kiemeney, L. A.; Kjaer, S. K.; Koushik, A.; Lambrechts, D.; Le, N. D.; Levine, D. A.; Massuger, L. F.; Matsuo, K.; May, T.; McNeish, I. A.; Menon, U.; Modugno, F.; Monteiro, A. N.; Moorman, P. G.; Moysich, K. B.; Ness, R. B.; Nevanlinna, H.; Olsson, H.; Onland-Moret, Nc.; Park, S. K.; Paul, J.; Pearce, C. L.; Pejovic, T.; Phelan, C. M.; Pike, M. C.; Ramus, S. J.; Riboli, E.; Rodriguez-Antona, C.; Romieu, I.; Sandler, D. P.; Schildkraut, J. M.; Setiawan, V. W.; Shan, K.; Siddiqui, N.; Sieh, W.; Stampfer, M. J.; Sutphen, R.; Swerdlow, A. J.; Szafron, L. M.; Teo, S. H.; Tworoger, S. S.; Tyrer, J. P.; Webb, P. M.; Wentzensen, N.; White, E.; Willett, W. C.; Wolk, A.; Woo, Y. L.; Wu, A. H.; Yan, L.; Yannoukakos, D.; Chenevix-Trench, G.; Sellers, T. A.; Pharoah, P. D.; Zheng, W.; Long, J. Genetic Data from Nearly 63,000 Women of European Descent Predicts DNA Methylation Biomarkers and Epithelial Ovarian Cancer Risk. Cancer research 2018, 79 (3), 505. 10.1158/0008-5472.CAN-18-2726.

(50) Li, T.; Fu, J.; Zeng, Z.; Cohen, D.; Li, J.; Chen, Q.; Li, B.; Liu, X. S. TIMER2.0 for Analysis of Tumor-Infiltrating Immune Cells. Nucleic Acids Research 2020, 48 (W1), W509. 10.1093/nar/gkaa407.

(51) Knecht, H.; Mai, S. LMP1 and Dynamic Progressive Telomere Dysfunction: A Major Culprit in EBV-Associated Hodgkin’s Lymphoma. Viruses 2017, 9 (7), 164. 10.3390/v9070164.

(52) Armanios, M.; Blackburn, E. H. The Telomere Syndromes. Nat Rev Genet 2012, 13 (10), 693–704. 10.1038/nrg3246.

(53) Lajoie, V.; Lemieux, B.; Sawan, B.; Lichtensztejn, D.; Lichtensztejn, Z.; Wellinger, R.; Mai, S.; Knecht, H. LMP1 Mediates Multinuclearity through Downregulation of Shelterin Proteins and Formation of Telomeric Aggregates. Blood 2015, 125 (13), 2101. 10.1182/blood-2014-08-594176.

(54) Cruz-Garcia, D.; Vazquez-Martinez, R.; Peinado, J. R.; Anouar, Y.; Tonon, M. C.; Vaudry, H.; Castaño, J. P.; Malagon, M. M. Identification and Characterization of Two Novel (Neuro)Endocrine Long Coiled-Coil Proteins. FEBS Letters 2007, 581 (17), 3149–3156. 10.1016/j.febslet.2007.06.002.

(55) Mu, X.; Liu, S.-J.; Zheng, L.-Y.; Ouyang, C.; Abdalla, A. M. E.; Wang, X.-X.; Chen, K.; Yang, F.-F.; Meng, N. The Long Coiled-Coil Protein NECC2 Regulates oxLDL-Induced Endothelial Oxidative Damage and Exacerbates Atherosclerosis Development in Apolipoprotein E −/− Mice. Free Radical Biology and Medicine 2024, 216, 106–117. 10.1016/j.freeradbiomed.2024.03.001.

(56) Ubel, C.; Mousset, S.; Trufa, D.; Sirbu, H.; Finotto, S. Establishing the Role of Tyrosine Kinase 2 in Cancer. Oncoimmunology 2013, 2 (1), e22840. 10.4161/onci.22840.

(57) Liu, X.; Sadaoka, T.; Krogmann, T.; Cohen, J. I. Epstein-Barr Virus (EBV) Tegument Protein BGLF2 Suppresses Type I Interferon Signaling To Promote EBV Reactivation. Journal of Virology 2020, 94 (11), e00258. 10.1128/JVI.00258-20.

(58) Geiger, T. R.; Martin, J. M. The Epstein-Barr Virus-Encoded LMP-1 Oncoprotein Negatively Affects Tyk2 Phosphorylation and Interferon Signaling in Human B Cells. J Virol 2006, 80 (23), 11638–11650. 10.1128/JVI.01570-06.

(59) Reed, J. C. Bcl-2 Family Proteins: Regulators of Apoptosis and Chemoresistance in Hematologic Malignancies. Semin Hematol 1997, 34 (4 Suppl 5), 9–19.

(60) Vatte, C.; Al-Amri, A. M.; Cyrus, C.; Chathoth, S.; Ahmad, A.; Alsayyah, A.; Al-Ali, A. Epstein-Barr Virus Infection Mediated TP53 and Bcl-2 Expression in Nasopharyngeal Carcinoma Pathogenesis. Molecular and Clinical Oncology 2021, 15 (6), 260. 10.3892/mco.2021.2422.

(61) Thai, A. A.; Young, R. J.; Bressel, M.; Kelly, G. L.; Sejic, N.; Tsao, S. W.; Trigos, A.; Rischin, D.; Solomon, B. J. Characterizing and Targeting of BCL-2 Family Members in Nasopharyngeal Carcinoma. Head Neck 2024. 10.1002/hed.27973.

(62) Pei, Y.; Wong, J. H. Y.; Robertson, E. S. Targeted Therapies for Epstein-Barr Virus-Associated Lymphomas. Cancers (Basel) 2020, 12 (9), 2565. 10.3390/cancers12092565.

(63) Wyżewski, Z.; Mielcarska, M. B.; Gregorczyk-Zboroch, K. P.; Myszka, A. Virus-Mediated Inhibition of Apoptosis in the Context of EBV-Associated Diseases: Molecular Mechanisms and Therapeutic Perspectives. International Journal of Molecular Sciences 2022, 23 (13), 7265. 10.3390/ijms23137265.

(64) Banerjee, S.; Uppal, T.; Strahan, R.; Dabral, P.; Verma, S. C. The Modulation of Apoptotic Pathways by Gammaherpesviruses. Front Microbiol 2016, 7, 585. 10.3389/fmicb.2016.00585.

(65) Hirokawa, N.; Noda, Y.; Okada, Y. Kinesin and Dynein Superfamily Proteins in Organelle Transport and Cell Division. Curr Opin Cell Biol 1998, 10 (1), 60–73. 10.1016/s0955-0674(98)80087-2.

(66) Exertier, P.; Javerzat, S.; Wang, B.; Franco, M.; Herbert, J.; Platonova, N.; Winandy, M.; Pujol, N.; Nivelles, O.; Ormenese, S.; Godard, V.; Becker, J.; Bicknell, R.; Pineau, R.; Wilting, J.; Bikfalvi, A.; Hagedorn, M. Impaired Angiogenesis and Tumor Development by Inhibition of the Mitotic Kinesin Eg5. Oncotarget 2013, 4 (12), 2302–2316. 10.18632/oncotarget.1490.

(67) Liu, S.-L.; Lin, H.-X.; Qiu, F.; Zhang, W.-J.; Niu, C.-H.; Wen, W.; Sun, X.-Q.; Ye, L.-P.; Wu, X.-Q.; Lin, C.-Y.; Song, L.-B.; Guo, L. Overexpression of Kinesin Family Member 20A Correlates with Disease Progression and Poor Prognosis in Human Nasopharyngeal Cancer: A Retrospective Analysis of 105 Patients. PLoS ONE 2017, 12 (1), e0169280. 10.1371/journal.pone.0169280.

(68) Tang, Y.; Nakada, M. T.; Rafferty, P.; Laraio, J.; McCabe, F. L.; Millar, H.; Cunningham, M.; Snyder, L. A.; Bugelski, P.; Yan, L. Regulation of Vascular Endothelial Growth Factor Expression by EMMPRIN via the PI3K-Akt Signaling Pathway. Molecular Cancer Research 2006, 4 (6), 371–377. 10.1158/1541-7786.MCR-06-0042.

(69) Krag, C.; Malmberg, E. K.; Salcini, A. E. PI3KC2α, a Class II PI3K, Is Required for Dynamin-Independent Internalization Pathways. Journal of Cell Science 2010, 123 (24), 4240–4250. 10.1242/jcs.071712.

(70) Gaidarov, I.; Smith, M. E.; Domin, J.; Keen, J. H. The Class II Phosphoinositide 3-Kinase C2alpha Is Activated by Clathrin and Regulates Clathrin-Mediated Membrane Trafficking. Mol Cell 2001, 7 (2), 443–449. 10.1016/s1097-2765(01)00191-5.

(71) Foster, F. M.; Traer, C. J.; Abraham, S. M.; Fry, M. J. The Phosphoinositide (PI) 3-Kinase Family. J Cell Sci 2003, 116 (Pt 15), 3037–3040. 10.1242/jcs.00609.

(72) Simabuco, F. M.; Morello, L. G.; Aragão, A. Z. B.; Paes Leme, A. F.; Zanchin, N. I. T. Proteomic Characterization of the Human FTSJ3 Preribosomal Complexes. J Proteome Res 2012, 11 (6), 3112–3126. 10.1021/pr201106n.

(73) Zhuang, Q.; Dai, Z.; Xu, X.; Bai, S.; Zhang, Y.; Zheng, Y.; Xing, X.; Hu, E.; Wang, Y.; Guo, W.; Zhao, B.; Zeng, Y.; Liu, X. RNA Methyltransferase FTSJ3 Regulates the Type I Interferon Pathway to Promote Hepatocellular Carcinoma Immune Evasion. Cancer Research 2024, 84 (3), 405–418. 10.1158/0008-5472.CAN-23-2049.

(74) Ringeard, M.; Marchand, V.; Decroly, E.; Motorin, Y.; Bennasser, Y. FTSJ3 Is an RNA 2’-O-Methyltransferase Recruited by HIV to Avoid Innate Immune Sensing. Nature 2019, 565 (7740), 500–504. 10.1038/s41586-018-0841-4.

(75) Tuna, M.; Amos, C. I.; Mills, G. B. Molecular Mechanisms and Pathobiology of Oncogenic Fusion Transcripts in Epithelial Tumors. Oncotarget 2019, 10 (21), 2095. 10.18632/oncotarget.26777.

(76) Shirakawa, R.; Horiuchi, H. Ral GTPases: Crucial Mediators of Exocytosis and Tumourigenesis. J Biochem 2015, 157 (5), 285–299. 10.1093/jb/mvv029.

(77) Yoshimachi, S.; Shirakawa, R.; Cao, M.; Trinh, D. A.; Gao, P.; Sakata, N.; Miyazaki, K.; Goto, K.; Miura, T.; Ariake, K.; Maeda, S.; Masuda, K.; Ishida, M.; Ohtsuka, H.; Unno, M.; Horiuchi, H. Ral GTPase–Activating Protein Regulates the Malignancy of Pancreatic Ductal Adenocarcinoma. Cancer Science 2021, 112 (8), 3064. 10.1111/cas.14970.

(78) Concepcion, A. R.; Wagner, L. E.; Zhu, J.; Tao, A. Y.; Yang, J.; Khodadadi-Jamayran, A.; Wang, Y.-H.; Liu, M.; Rose, R. E.; Jones, D. R.; Coetzee, W. A.; Yule, D. I.; Feske, S. The Volume-Regulated Anion Channel LRRC8C Suppresses T Cell Function by Regulating Cyclic Dinucleotide Transport and STING-P53 Signaling. Nat Immunol 2022, 23 (2), 287–302. 10.1038/s41590-021-01105-x.

(79) Manolios, N.; Papaemmanouil, J.; Adams, D. J. The Role of Ion Channels in T Cell Function and Disease. Frontiers in Immunology 2023, 14, 1238171. 10.3389/fimmu.2023.1238171.

(80) Yuan, X. H.; Li, Y.; Li, P. Regulatory Effects of Linc00486 on Biological Characteristics of Breast Cancer Cells. Revista argentina de clínica psicológica 2020, 29 (3), 356–363.

(81) Lin, W.; Yip, Y. L.; Jia, L.; Deng, W.; Zheng, H.; Dai, W.; Ko, J. M. Y.; Lo, K. W.; Chung, G. T. Y.; Yip, K. Y.; Lee, S.-D.; Kwan, J. S.-H.; Zhang, J.; Liu, T.; Chan, J. Y.-W.; Kwong, D. L.-W.; Lee, V. H.-F.; Nicholls, J. M.; Busson, P.; Liu, X.; Chiang, A. K. S.; Hui, K. F.; Kwok, H.; Cheung, S. T.; Cheung, Y. C.; Chan, C. K.; Li, B.; Cheung, A. L.-M.; Hau, P. M.; Zhou, Y.; Tsang, C. M.; Middeldorp, J.; Chen, H.; Lung, M. L.; Tsao, S. W. Establishment and Characterization of New Tumor Xenografts and Cancer Cell Lines from EBV-Positive Nasopharyngeal Carcinoma. Nat Commun 2018, 9 (1), 4663. 10.1038/s41467-018-06889-5.

(82) Yip, Y. L.; Lin, W.; Deng, W.; Jia, L.; Lo, K. W.; Busson, P.; Vérillaud, B.; Liu, X.; Tsang, C. M.; Lung, M. L.; Tsao, S. W. Establishment of a Nasopharyngeal Carcinoma Cell Line Capable of Undergoing Lytic Epstein-Barr Virus Reactivation. Lab Invest 2018, 98 (8), 1093–1104. 10.1038/s41374-018-0034-7.

(83) Cheung, S. T.; Huang, D. P.; Hui, A. B.; Lo, K. W.; Ko, C. W.; Tsang, Y. S.; Wong, N.; Whitney, B. M.; Lee, J. C. Nasopharyngeal Carcinoma Cell Line (C666-1) Consistently Harbouring Epstein-Barr Virus. Int J Cancer 1999, 83 (1), 121–126. https://doi.org/10.1002/(sici)1097-0215(19990924)83:1<121::aid-ijc21>3.0.co;2-f.

